# Hub architecture of the human structural connectome: Links to aging and processing speed

**DOI:** 10.1101/2023.01.28.525980

**Authors:** Xin Li, Alireza Salami, Jonas Persson

**Affiliations:** Aging Research Center, Karolinska Institute and Stockholm University, 171 65, Stockholm, Sweden; Umeå Center for Functional Brain Imaging (UFBI), Umeå University, 901 87, Umeå, Sweden; Wallenberg Centre for Molecular Medicine, Umeå University, 901 87, Umeå, Sweden; Department of Integrative Medical Biology, Umeå University, 901 87, Umeå, Sweden; Center for Lifespan Developmental Research (LEADER), School of Behavioral, Social and Legal Sciences, Örebro University, 701 82, Örebro, Sweden

**Keywords:** connectome, fiber bundle capacity, processing speed, diffusion-weighted imaging, aging

## Abstract

The human structural brain network, or connectome, has a rich-club organization with a small number of brain regions showing high network connectivity, called hubs. Hubs are centrally located in the network, energy costly, and critical for human cognition. Aging has been associated with changes in brain structure, function, and cognitive decline, such as processing speed. At a molecular level, the aging process is a progressive accumulation of oxidative damage, which leads to subsequent energy depletion in the neuron and causes cell death. However, it is still unclear how age affects hub connections in the human connectome. The current study aims to address this research gap by utilizing a novel measure of structural connectivity strength, fiber bundle capacity (FBC), which is derived from Constrained Spherical Deconvolution (CSD) modeling of white-matter fiber bundles. FBC represents the capacity of a fiber bundle to transfer information and is a less biased measure for quantifying connection strength within biological pathways. We found that hubs exhibit longer-distance connections and higher metabolic rates compared to peripheral brain regions, suggesting that hubs are biologically costly. Although the landscape of structural hubs was relatively age-invariant, there were wide-spread age effects on FBC in the connectome. Critically, these age effects were larger in connections within hub compared to peripheral brain connections. These findings were supported by both a cross-sectional sample with wide age-range (N=137) and a longitudinal sample across 5 years (N=83). Moreover, our results demonstrated that associations between FBC and processing speed were more concentrated in hub connections than chance level, and FBC in hub connections mediated the age-effects on processing speed. Overall, our findings indicate that structural connections of hubs, which demonstrate greater energy demands, are particular vulnerable to aging. The vulnerability may contribute to age-related impairments in processing speed among older adults.

## Introduction

The human brain can be viewed as a network at the macroscopic level, with individual brain regions, nodes, interconnected by white matter (WM) fibers. The structural brain network is a complex scale-free network (Power et al., 2013; van den Heuvel and Sporns, 2011), which is characterized by the probability of nodes degree (i.e., the number of connections to other nodes) in the network decaying as a power law, free of a characteristic scale, for large degrees. A scale-free network indicates the existence of a subset of nodes, referred to as hubs, having a large proportion of connections. The spatial distance among hubs is relatively longer compared to peripheral nodes (Alexander-Bloch et al., 2013; Van Den Heuvel et al., 2012). This indicates that brain hubs have high anatomical wiring cost and are critical for integrating brain sub-networks and information transformation among long-distance brain regions. In addition, hubs, especially in cortical areas, have greater metabolic rate (Liang et al., 2013; Tomasi et al., 2013; Vaishnavi et al., 2010), suggesting that maintaining normal function of hubs is energy costly. Hubs are disproportionately expanded in humans than other primates (Hill et al., 2010). Thus, the existence of hubs might partly explain cognitive capabilities in humans (Buckner and Krienen, 2013). Brain lesions linked to various brain disorders, including schizophrenia and Alzheimer’s disease, were concentrated in hubs compared to peripheral brain regions (Buckner et al., 2009; Crossley et al., 2014). Possibly, the high metabolic cost and long-distance connections of hubs make them vulnerable to pathological damage.

Aging is associated with changes in WM integrity/fiber densities (Kelley et al., 2021; Sexton et al., 2014). A recent longitudinal study demonstrated structural connectome reorganization as indicated by decreased integration and increased segregation over 5-years (Coelho et al., 2021). The underlying reason for this structural reorganization is unknown. At a molecular level, aging is characterized by a progressive accumulation of damage induced by reactive oxygen, nitrogen, and carbonyl species (RONCS). RONCS is the by-products of various catabolic and anabolic processes that can lead to subsequent energy depletion in the neurons and cause cell death. Based on this chain of events, one might hypothesize that long-distance hub connections are more affected by decreased energy metabolism induced by the oxidative damage during normal aging. However, no study has systematically examined if connections within hubs are more effected by age than non-hubs in the structural connectome.

Moreover, previous diffusion tensor imaging (DTI) studies have shown that WM integrity can partly explain age-related differences in tasks that require speeded responses (Andersson et al., 2022; Li et al., 2022; Salami et al., 2012). Indeed, processing speed has been correlated with higher-order cognition, such as working memory and fluid intelligence (Ackerman et al., 2002; Waters and Caplan, 2005). Lower processing speed may lead to accelerated cognitive impairment across task domains in older adults (Finkel et al., 2005). It is currently unknown whether hub connections are more important for cognitive performance, such as processing speed, in older adults.

In this study, we investigate the age effects on structural hubs and its relationship with processing speed using network-based and graph-theoretical approaches (Figure 1). The distinction between hubs and non-hubs in a brain network is relatively arbitrary. Thus, our graph-theoretical analysis was conducted on varying thresholds to define hubs instead of on a hard cut-off. Both cross-sectional data with a wide age range from 25-85 years and longitudinal data with a five-year interval were used in the study. We hypothesize that, (1) the *locations* of hubs in the structural brain network remain stable across the adults life-span, but that (2) there are wide-spread age effects on *connectivity strength* across the connectome; more importantly (3) the age effects are larger within hub compared to peripheral connections; and finally that (4) the connectivity strength in hub connections are more strongly related to processing speed, such that hub connections can explain processing speed impairments in older adults.

**Figure 1.**
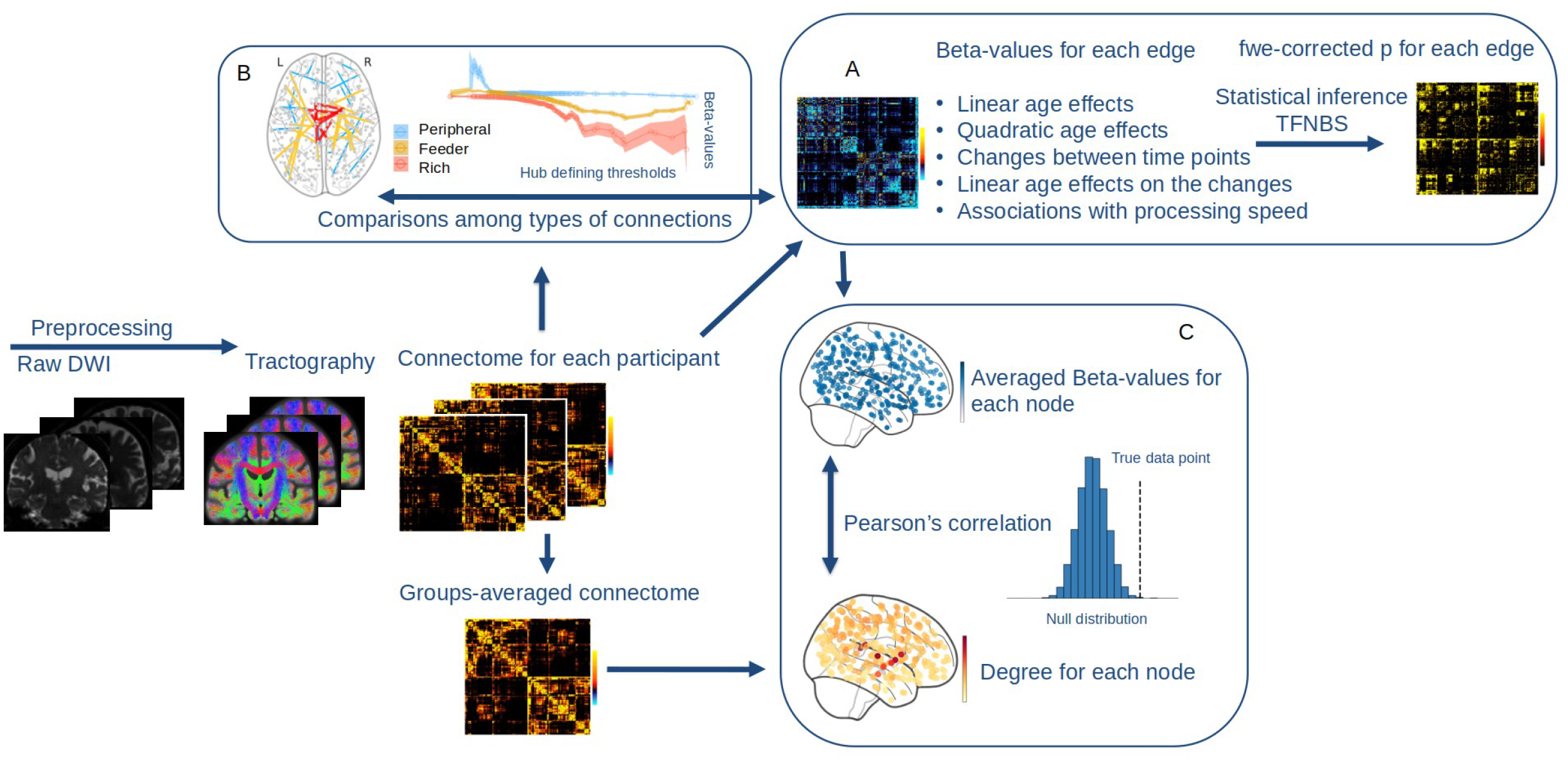
Schematic illustration of the analytical pipeline. After preprocessing, (A) connectivity strength in the connectome was associated with age or speed using regression models. Threshold-free network-based analysis (TFNBS) was applied to estimate the beta- and p-values for each edge. (B) Degree/hubness of each node was derived from the connectome using graph theory. Rich/feeder/peripheral connections were defined based on multiple thresholds of degree/hubness. The beta-values for each edge were compared among rich/feeder/peripheral connections. (C) Beta-values were averaged for each of the nodes, which were subsequently correlated with the map of degree of each node.

## Methods

### 1. Participants

The participants in the current study are part of the Betula prospective cohort study on memory, health and aging (Nilsson et al., 1997; Nyberg et al., 2020). The Betula study was approved by the Regional Ethical Review Board in Umeå, and all participants gave written informed consent. The Betula project was launched in 1988 and 7 waves of health and neuropsychology data until 2017 (T1-T7) have been collected. The time interval between the waves is five years. The current study is based on the neuroimaging subsample recruited from the T5 to T7 waves. The original sample size of the 3 waves is 376 (T5), 230 (T6), and 103 (T7). We excluded participants with dementia, stroke, epilepsy, Parkinson’s disease, multiple sclerosis, hydrocephalus, head surgery, deviant brain morphology, vascular brain lesions and infarcts. Dementia was assessed at baseline and reassessed for each wave using a three-step procedure, according to the Diagnostic and Statistical Manual of Mental Disorders, 4th edition (DSM–IV; American Psychiatric Association, 1994). First, an overall evaluation was performed by an examining physician. The diagnosis was then compared with a second independent diagnosis based on scores from several cognitive tests. In cases of disagreement, a supervising physician made a third and final diagnosis. Cross-sectional analysis was conducted on the data of wave T5, which has the largest sample size. In the age-comparison analyses, five age groups: 25-45, 50-55, 60-65, 70-75 and 80-85 years were defined and compared. In this sample, echo time (TE) varies within waves, and between T5 and T6/T7 waves. Since TE may potentially influence the estimated WM fiber densities, longitudinal analyses were carried out for T6 and T7 only. Participants with different TEs at T5 and with changes of TE between T6 and T7 were excluded. The demographic information and behavioral performance of processing speed of the final sample are present in Table 1.

**Table 1.** Demographic information and cognitive performance

|  | T5 | T6 | T7 |
| --- | --- | --- | --- |
|  | Cross-sectional design | Longitudinal design |  |
| N | 137 | 84 |  |
| Age | 62 (25-85) | 69.2 (55-80) at T6 |  |
| Sex | 100/37 | 42/42 |  |
| Education | 13 | 13.9 |  |
| MMSE | 28.1 (24-30) | 28.2 (24-30) | 28.1 (24-30) |
| Processing speed | 55.4 | 50.6 | 50.3 |

### 2. Imaging acquisition and processing

Diffusion weighted imaging (DWI) data were collected on a 3T Discovery MR750 (General Electric) scanner with a 32-channel head coil. Data were acquired using a single-shot, spin-echo-planar, T2-weighted sequence, with a spatial resolution of 0.98 × 0.98 × 2 mm. The sequence parameters were: TR = 8.0 s, 64 slices with no gap in between, 256 × 256 matrix (FOV = 250 mm), 90° flip angle, b = 1000 s/mm2 and six b = 0 images. At baseline and the first follow up, 3 DWI sessions were included for each participant. At the second follow up, only one session was included in the protocol. To retain consistency in data analyses across multiple time point (T5-T7), only the first scanning session (for baseline and first follow up) were included in the analysis. TE is 84.4 ms at T5 and varied between 81.2 ms to 82.8 ms at T6/T7. All participants were examined on the same scanner with no software or hardware update during the data-collection period.

The raw DWI data was first pre-processed using MRtrix 3.0, including denoising, unringing and motion correction, topup, eddy current correction, and bias field correction. We used Synb0 (Schilling et al., 2020) to correct for the spatial distortions caused by susceptibility induced off-resonance fields. This approach synthesizes an undistorted b0 image based on a distorted b0 and a T1 image using deep learning. The ventricles, scalp and nonparenchymal regions within the images were removed using the BET tool as implemented in FSL (Smith, 2002).

We used the constrained spherical deconvolution (CSD; Tournier et al., 2004, 2007) to obtain the Fiber Orientation Distributions (FODs) in each voxel. This was done by first estimating the response functions for single-fiber WM as well as gray matter (GM) and CSF using an unsupervised method (Dhollander et al., 2019). Single-Shell 3-Tissue CSD (SS3T-CSD) was performed to obtain WM-like FODs as well as GM-like and CSF-like compartments in all voxels (Dhollander and Connelly, 2016), using MRtrix3Tissue (https://3Tissue.github.io), a fork of MRtrix3 (Tournier et al., 2019). WM FOD images were then corrected for intensity inhomogeneities and used in subsequent analyses.

Whole-brain Anatomically Constrained Tractography (ACT, Smith et al., 2012) was performed on the WM FOD images. ACT is an additional module to the streamlines tractography, which uses T1 images as anatomical priors to constrain the streamlines beginning, traversing, and ending in anatomically plausible regions. This was achieved by processing and segmenting the T1-image into 5 different tissue types, cortical GM, sub-cortical GM, WM, CSF, and pathological tissue. The segmented images were aligned to the B0 images for each individual using ANTs and were then fed into the whole-brain tractography as a mask. ACT improves the biological accuracy of tractography, and greatly decrease false positive streamlines. Ten million tractography streamlines were reconstructed, with the GM–WM interface used as seed regions. The generated tractograms were filtered using Spherical-deconvolution informed filtering of tracks (SIFT2, Smith et al., 2015).

### 3. Connectome construction

Structural connectomes were generated based on tractograms and the atlas-based parcellation images. The network includes 179 atlas regions/nodes on each cortical hemisphere, based on HCPMMP 1.0, with an additional 16 subcortical regions based on the FreeSurfer segmentation. Two regions in the temporal cortex spans only 2 voxels with no associated streamlines and were therefore excluded in the analysis. The cerebellum and the brainstem were not included in some images because of restricted field of view and these regions were therefore omitted from further analyses. A novel measurement, fiber bundle capacity (FBC, Smith et al., 2020), was used to estimate the structural connectivity strength/edges of the network. FBC represents the capacity of a fiber bundle to transfer information. Compared to the commonly used connectivity measures, such as number of streamlines, FBC is reflective of the underlying biological connectivity, i.e., fiber density (Smith et al., 2020), and is less influenced by brain volume or streamline length. Moreover, although the number of streamlines generated in tractography is fixed for each participant, and the volumes of each parcellated brain region is adjusted during the post-analysis, the number of streamlines assigned to a particular pair of parcels might still be biased due to many factors that cannot be controlled for (Smith et al., 2020). Thus, we consider FBC to be a more robust and less biased measure.

Since the distribution of FBC is heavily right-skewed (Figure S1), we used the log-transformed FBC to quantify the connection weights of the network (i.e., network edges) in the analysis. To further minimize false-positive connections, we used two criteria to select connections to be included in subsequent analyses (Arnatkeviciute et al., 2021) (i) connections that are present in at least 30 % of participants; and (ii) amongst the τ % strongest edges to achieve the desired network density. The densities (proportion of non-zero connections) of the mean connectome were 64.4 %, 64.7 %, 68.5 %, 67.4 %, and 57.8 % for age groups. After thresholding (i), the densities of the mean connectome were 46.4 % for all age groups younger than 80 years, and 46.2% for 80-85 group. The density distributions for each participant are shown in Figure S2. Weak connections might introduce randomness that could influence graph-theoretical measurements (van den Heuvel et al., 2017). We therefore examined our main results across a range of densities (τ = 5 %, 10 %, 15 %, 20 %, 25 %, 30%), and on the mean connectome based on these six densities. This ensured that a proportion of weak/false-positive connections were further removed for all the participants after thresholding (ii). Moreover, after thresholding (ii), the density distributions among age groups were different for τ = 15%, 20 %, 25 %, 30% (p-values < 0.05), and were comparable for τ = 5 %, 10 %. Thus, the selection of these thresholds also aims to test whether our findings are affected by the age effects on network densities. The analysis using τ = 30 % were presented in the main results. The replication using τ = 5 %, 10 %, 15 %, 20 %, 25 %, and the mean connectome are reported as supplementary results.

### 4. Characteristics of the human connectome

To characterize the topographical organization of the connectome, we calculated 3 metrics, degree, rich-club coefficient, and connection distance on the averaged connectome of the baseline sample. Degree was computed for each node as the total number of edges in the network connected to this node. At a particular degree threshold *k*, hubs are defined as the nodes with degree > *k*, and non-hubs are the nodes with degree <= *k*. Connections are subsequently labeled as rich (hub–hub), feeder (hub–non-hub), and peripheral (non-hub–non-hub) connections. Rich-club coefficient measures the extent to which hub regions in a network connect between each other.

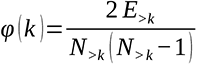

where *N_k_* is the number the nodes with degree > *k*, and *E_k_* are the number of edges among nodes with degree > *k* (Colizza et al., 2006). Since φ(k) is expected to be larger within nodes with higher degree, φ(*k*) can be normalized by dividing φ(k) of the random networks. We generated 1000 random null networks while retaining the same network degree, φ_random_(k) was then computed for

each network and averaged.

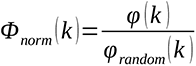

Φ_norm_> 1 indicates a rich-club organization. Statistical significance of Φ_norm_ was tested based on the distribution of the φ_random_(k). Degree and rich-club coefficients were calculated using the Brain Connectivity Toolbox. Euclidean distance is defined as the straight-line distance between the centroids of pair of regions. This is a commonly used measure of wiring cost of connections.

In addition to defining hubs using degree centrality, an unweighted measurement of connectivity strength, hubs are defined using “hubness”, which is based on a sum of several graph metrics, including weighted connectivity strength, path length, weighted betweenness centrality, and weighted clustering-coefficient (Van Den Heuvel et al., 2010). We explored the distribution of “hubness” across regions of the structural brain networks, and also addressed the question if “hubness” resulted in identification of the same regions as those being hubs. To calculate the hubness of each node, we sum the rank of connectivity strength and betweenness centrality in an ascending order, and path length and clustering-coefficient in a descending order. The path length of a node refers to the averaged minimal distance between this node and all the nodes in the network. The equation is:

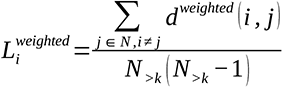

with

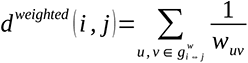

where 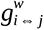 is the shortest path between node *i* and *j, and w_uv_* is the connection weights (i.e., log-transformed FBC) of all pairs of nodes *u* and *v* on the shorted distance 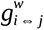. Global efficiency is the average inverse shortest path length in the network. Betweenness centrality of a node is defined as the proportion of the shortest paths that passed through this node. The equation is:

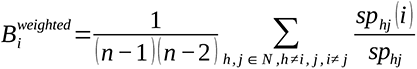

where *sp_hj_* (*i*) is the number of shortest paths in the network that passes through node *i*.

Clustering-coefficient of a node represents the probability that the neighbors of this node are also interconnected.

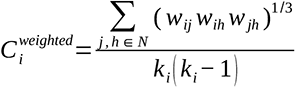

where *k_i_* is number of nodes that are connected to node i, and 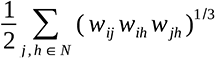 is the geometric mean of triangles around *i*.

We further investigated whether connectome hubs had higher metabolic rate compared to non-hubs using the brain regional metabolism data provided by a separate positron emission tomography study (Vaishnavi et al., 2010). In this particular study, several cerebral metabolic measurements, including glycolytic index (GI), cerebral metabolic rate for oxygen (CMRO2), glucose (CMRGlu) and the cerebral blood flow (CBF), were acquired in 33 normal adults during rest. The values of the measurements were then assigned to 82 Brodmann regions in the brain. We related mean connectome degrees to the metabolic measures across regions using Pearson’s correlation. Subcortical regions and left and right Brodmann area 26 were excluded from the analysis. Permutation was used to test statistical significance of the correlation against null models. We generated 1000 randomly shuffled brain maps (surrogate maps) with preserved similar spatial autocorrelation (SA) using BrainSMASH (Burt et al., 2020). Pearson correlations were performed between brain map representing the degree and 10.000 randomly shuffled but spatially constrained brain maps. P-values (P_permutation_) smaller than 0.05 were considered as being significant.

### 5. The influence of age on the connectome

#### 5.1 Threshold-free network-based analysis

For the cross-sectional analysis, we examined the effect of age on whole brain log-transformed FBC at the edge level (Equation1: log (*FBC*) ∼ *Age*+ *Sex*) using non-parametric permutation testing with the threshold-free network-based statistics (TFNBS, Vinokur et al., 2015) algorithm. This algorithm uses the concept of Threshold Free Cluster Enhancement (TFCE, Smith and Nichols, 2009) to the network domain, and outputs an enhanced matrix based on local cluster extent and magnitude. The enhanced matrix can then be turned into edge-wise p-values using permutation testing. TFNBS is more sensitive than Network Based Statistic (Zalesky et al., 2010), and does not require a hard cluster-forming threshold. Permutation was performed with n = 5000 data shuffling for statistics inference. We then added an age square term in the equation to investigate the quadratic age effects (Equation2: log (*FBC*) ∼ *Age*+ *Age*^2^+ *Sex*). TFNBS was performed in MRtrix3.0. For the longitudinal analysis, we examined changes in structural connectivity from T6 to T7 and the linear age effects on the changes in older adults across 5 years (Equation3: log (△ *FBC*) ∼ *Age*+*Sex*). P-values (P_TFNBS_) smaller than 0.05 were considered as being significant.

#### 5.2 Graph theoretical analyses

We compared the degrees of the connectomes between the five age groups (25-45, 50-55, 60-65, 70-75 and 80-85 years), and across time points from T6 to T7. Moreover, we tested the age effect on the overall nodal distribution of degree. This was done by first choosing a degree distribution of the averaged connectome of the 25-45 age bin as a youth-like degree organization, and then calculating the similarity between degree distribution for each participant and the youth-like degree organization using Pearson’s correlation. Linear and quadratic age effects on the similarity (correlation coefficients) were estimated using a linear model for all participants from 50 years old. To characterize whether the effect of age on connectivity strength is preferentially stronger in certain types of connections in the human brain, we used linear regression (Equations 1 and 2) to estimate the beta-values of linear and quadratic age effects on mean connectivity of rich, peripheral, and feeder connections across different hub-defined thresholds. Sex was included as a covariate in the analyses. For each type of connection under a specific threshold *k*, mean connectivity is represented as the sum of log-transformed FBC divided by the number of non-zero connections. Thus, the analysis will not be confounded by the fact that the number of connections was larger for higher degree nodes. Another way of examining the data is to examine whether the edge-wise beta-values are more concentrated in certain types of connections. This was done by comparing beta-values at the edge level among rich, peripheral, and feeder connections across hub-defined thresholds using Welch’s t test.

We then related the degree of the connectome to the beta-values of age-effects across nodes to further investigate whether the effects of age are larger in nodes with higher degrees. The output from TFNBS was a 374 × 374 matrix of beta-values of linear (or quadratic) age effects, which were turned into a vector of 374 mean beta-values by row-wise sum of the matrix divided by the number of non-zero connections. The vector of mean beta-values represents to what extent age influences the average connectivity strength from each node to the rest of the brain. This was then correlated with the degrees of each node using Pearson’s correlation. Statistical inferences were based on permutation tests with 10.000 SA-preserved surrogate maps.

For the longitudinal analysis, we examined whether the changes and the linear age effects on the changes of connectivity strength (Equation 3) were larger in rich compared to peripheral, or feeder connections. Similar to the cross-sectional analysis, this was done by (1) comparing the longitudinal changes or the age effects on the changes among rich, peripheral, and feeder connections across different hub-defining thresholds; (2) relating the degree of the connectome to the beta-values across nodes. Statistical inferences were based on permutation tests with 10.000 SA-preserved surrogate maps.

In addition to degree, we also explored how other topological properties of the connectomes differed between the five age groups, and whether they change from T6 to T7. Several commonly used graph metrics were tested, including path length, global efficiency, betweenness centrality, and clustering coefficient.

### 6. The relation between structural connectivity and processing speed

The associations between whole brain log-transformed FBC and processing speed were examined using TFNBS controlling for age and sex (Equation 3, *Speed* ∼ *Age*+ *Sex*+log (*FBC*)). To test whether connections that were linked to hubs are critical for processing speed, we compared the median of the degrees for the regions that demonstrated significant connections associated with speed in the TFNBS analysis with median degree of the rest of the brain. In addition, we investigated whether the connections that were potentially related to speed from TFNBS were more concentrated in rich, peripheral, or feeder connections. This was examined by first identifying hubs/non-hubs based on an arbitrary nodal degree threshold (top 20% of the degree distribution). The averaged beta-values of the associations with speed were compared among rich, peripheral and feeder connections. The statistical inferences were made based on SA-preserved permutation (Burt et al., 2020).

We investigated whether structural connectivity mediates the relation between age and processing speed. Averaged connectivity strength was extracted for the connections that were significantly associated with speed. We estimated the total effect of age on speed, the effect of age and averaged connectivity strength on speed, and the indirect mediating effect of age on speed via connectivity strength. The confidence interval of the mediation effect was produced using a quasi-Bayesian approximation using the “mediation” package in R (Tingley et al., 2014).

### 7. Replication and validation

To determine whether our findings are robust, we replicated the analyses using multiple alternative approaches and parameters. First, in our main analysis, we explore whether our main results, the heterogeneous age effects on rich/peripheral/feeder connections, are robust and replicable based on “hubness” (Van Den Heuvel et al., 2010). Second, the size of individual brain regions can influence the results. We therefore repeated the correlation analysis between the brain maps of degree and the age effects while controlling for regional brain volume. In addition, we replicated out main findings across different connectome density thresholding (τ = 5 %, 10 %, 15 %, 20 %, 25% and the mean connectome based on six density thresholds).

## Results

### 1. Characteristics of the group-level structural network

We found that the averaged structural network has a long-tailed degree distribution (Figure 2A), and the tail of the degree distribution (*k* > 115) follows a power-law (bootstrapped-p > 0.05, Clauset et al., 2009). This suggests that the network is a scale-free network with several hub nodes having high degrees. The nodes with 1 SD above the mean degree (around top 9% highest degree) include the following regions: the bilateral retrosplenial cortex, bilateral thalamus, bilateral putamen, bilateral caudate nucleus, bilateral globus pallidus, bilateral ventral diencephalon, bilateral hippocampus, two regions in the anterior cingulate cortex, one region in the left mid-cingulate, and one region in the left posterior cingulate (Figure 2B). φnorms within these hubs are significantly stronger than expected by chance and increase with the hub-defining threshold *k* increasing (Figure 2C). This suggests that the network connections demonstrate a rich-club organization, and hubs are more densely connected than chance level. The mean connection distance (a measure of wiring cost) increases as a function of increased hub-defining threshold *k*, and the mean distance among hubs are significantly larger (one-sided Welch’s t-test, p < 0.05) than the mean distance of peripheral and feeder connections across multiple thresholds (Figure 2D, black dots). In addition, across 79 Brodmann areas, mean degrees were correlated with several measures of regional brain metabolisms, including GI, CMRO_2_, CMRGlu, and CBF (Figure 2E). The results indicate greater metabolic rates in higher-degree compared to lower-degree brain areas across cortical regions.

**Figure 2.**
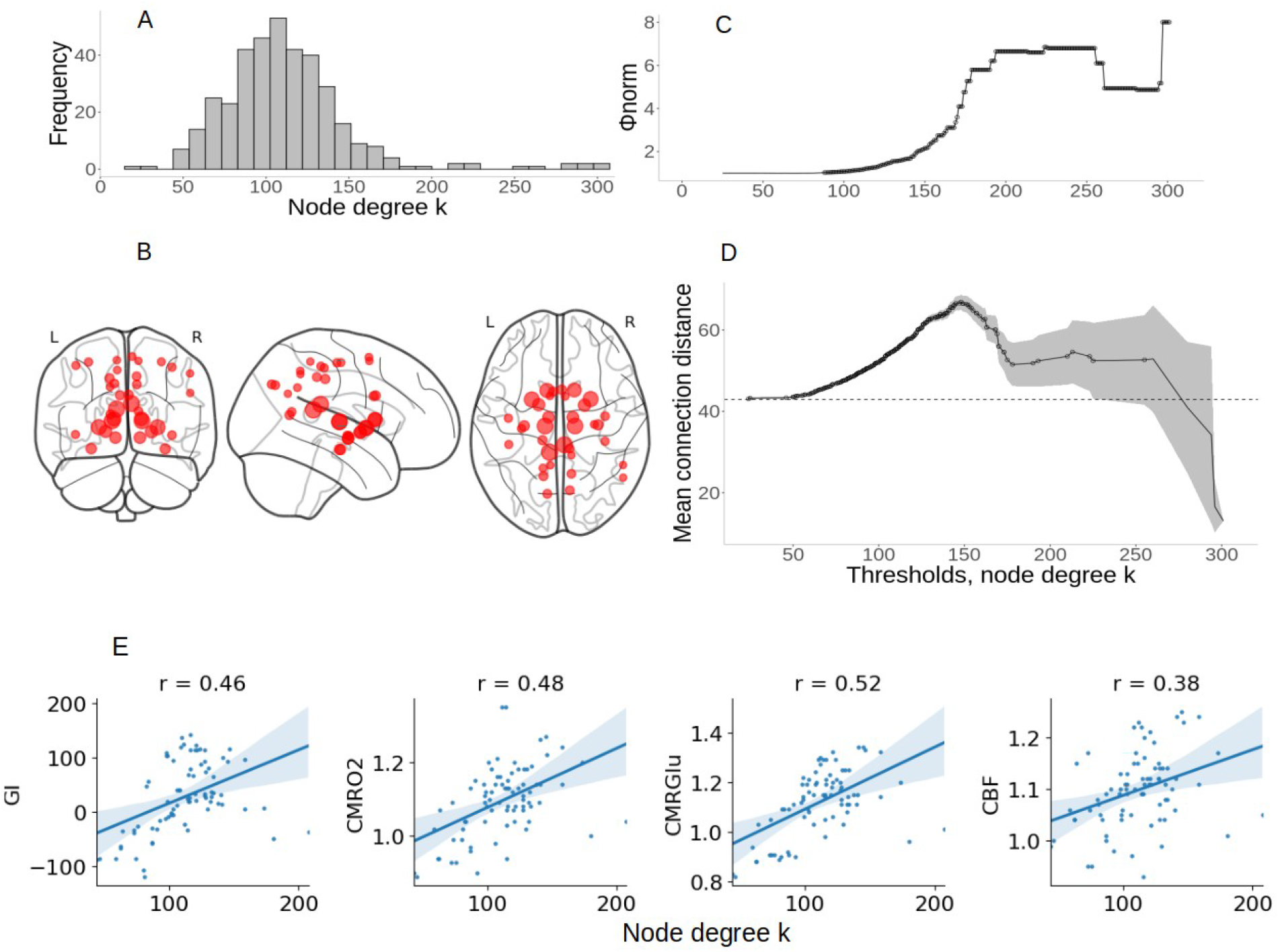
Network characteristics. (A) the long-tailed degree distribution across nodes indicates the presence of a small number of highly connected hubs. (B) The anatomical location of hubs defined by 1 SD higher than mean degree; the size indicated the degree of each node. (C) The normalized rich-club coefficient φ norm increases as a function of degree threshold (*k*) used to define hubs. Black circles indicate significant (P_permutation_ < 0.05) φnorm compared to 1000 degree-retained random networks. (D) Mean connection distance among rich club (hub–hub) connections increases as a function of degree threshold (*k*) used to define hubs. The shaded area depicts two standard errors around the mean. Black circles indicate significant larger distance among rich club connections compared to averaged feeder and peripheral connections. The dashed line shows the average connection distance across all network links. (E) Mean degrees are correlated with glycolytic index (GI, Pearson’s r(77) = 0.46, P_permutation_ < 0.05), the cerebral metabolic rate for oxygen (CMRO2, Pearson’s r(77) = 0.48, P_permutation_ < 0.05), cerebral metabolic rate for glucose (CMRGlu, Pearson’s r(77) = 0.52, P_permutation_ < 0.05), and the cerebral blood flow (CBF, Pearson’s r(77) = 0.38, P_permutation_ < 0.05). The statistical significances of Pearson’s correlation coefficients were based on 1000 permutation tests.

Degree was highly correlated with hubness, and with the other three measures of centrality, i.e., path length, betweenness centrality, and clustering-coefficient (Figure S3A). Hubs defined by degree are also consistently ranked for the measure of hubness (Figure S3B). The nodes with top 1 SD above mean hubness are as follows: the bilateral retrosplenium cortex, bilateral hippocampus, bilateral pallidum, bilateral ventral diencephalon, three regions in the bilateral posterior cingulate, bilateral thalamus, bilateral amygdala, and bilateral mid-cingulate. These regions are fully covered by the hubs defined using nodal degree. However, caudate and putamen were defined as hubs according to their high degree but were not hubsusing “hubness” as a definition, due to their relatively low betweenness centrality.

### 2. The influence of age on the brain connectome

Degrees were highly correlated and the identified hubs (brain regions with 1 SD higher than the mean degree) were similar between age groups and across the five-year time interval, suggesting relative stable spatial distributions of hubs in healthy adults of different ages. However, there was a significant age effect on the youth-like degree distributions, indicating that older participants have less youth-like degree distribution (p < 0.001). The distributions of the degrees between pairs of age bins/time points are presented in Figure S4. Anatomical locations of hubs for the five age groups are presented in Figure 3.

**Figure 3.**
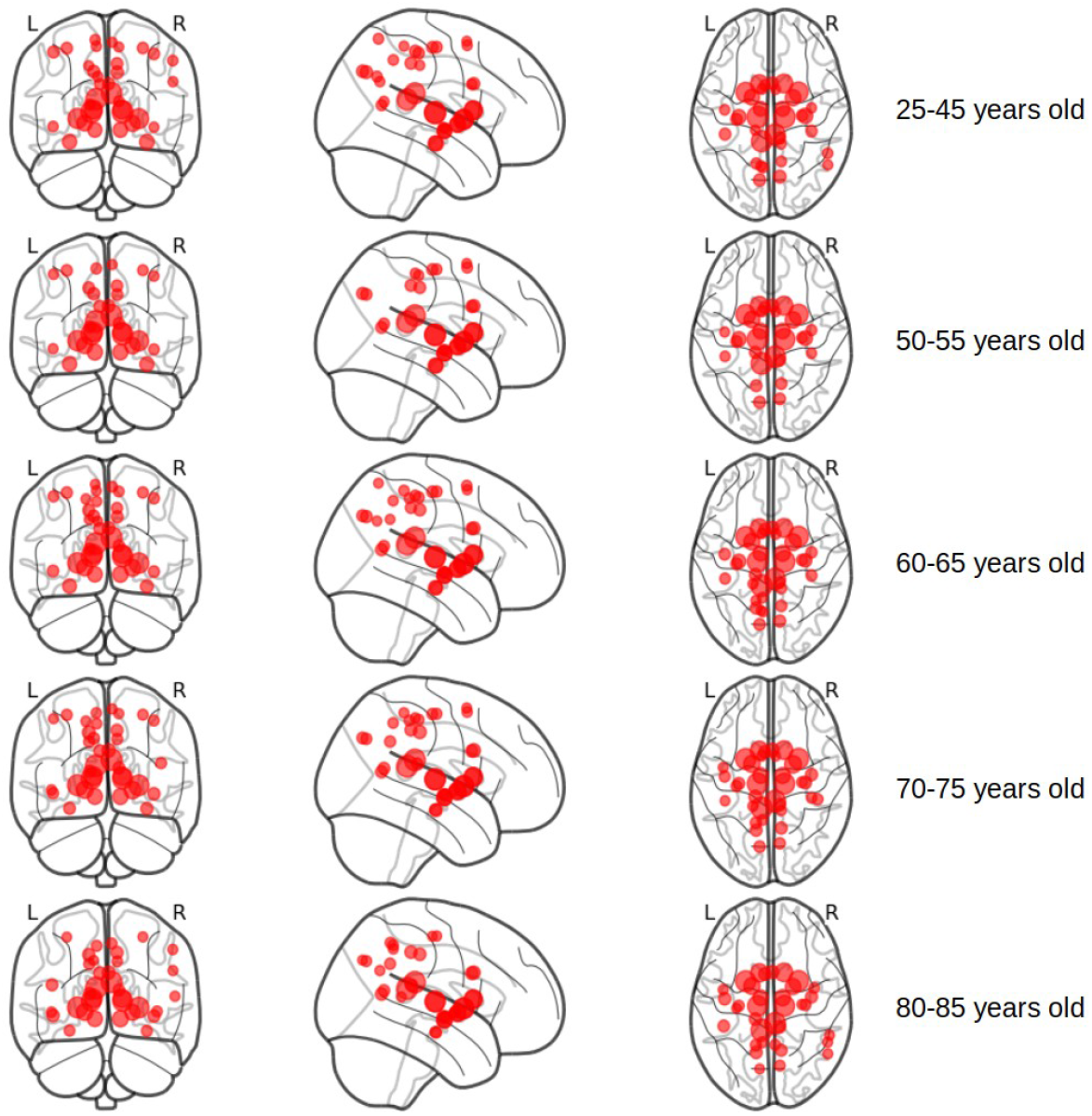
The anatomical locations of hubs that defined by 1 SD higher than mean degree across age groups; the size indicates the degree of each node.

#### 2.1 Age-related alterations of the structural connectome across the adult lifespan

TFNBS analyses showed both significant linear (Figure 4A) and quadratic (Figure S5A) age effects on multiple cortical–cortical connections and most of the cortical–subcortical connections, suggesting that older adults had globally lower structural connectivity strength compared to younger adults. More importantly, our results showed that both of the linear (Figure 4B) and quadratic (Figure S5B) age effects were largest for hub connections compared to peripheral, and feeder connections regardless of the thresholds used to define hubs. The linear age effects in hub connections with thresholds *k* from 24 to 260, and feeder connections with *k* from 51 to 301 are significantly larger than the peripheral connections. The quadratic age effects in hub connections with thresholds *k* from 46 to 280, and feeder connections with *k* from 51 to 301 are significantly larger than the peripheral connections. Mean linear/quadratic beta values were also larger for rich connections compared to feeder (linear: P < 0.005 for 175 > *k* > 46; quadratic: P < 0.005 for 168 > *k* > 85) and peripheral connections (linear: P < 0.0001 for *k* > 102; P < 0.005 for *k* > 103) across hub-defining thresholds (Figure S6). This suggests that higher negative beta-values are mostly concentrated in rich connections, compared to feeder or peripheral connections. In addition, we found significant correlation between the degrees and the averaged beta-values of the age effects (P_permutation_ < 0.0001, r = -0.496 for linear age effects; P_permutation_ < 0.05, r = -0.352 for quadratic age effects), indicating that brain regions with higher degree also showed more pronounced age-related decreases in their connectivity strength.

**Figure 4.**
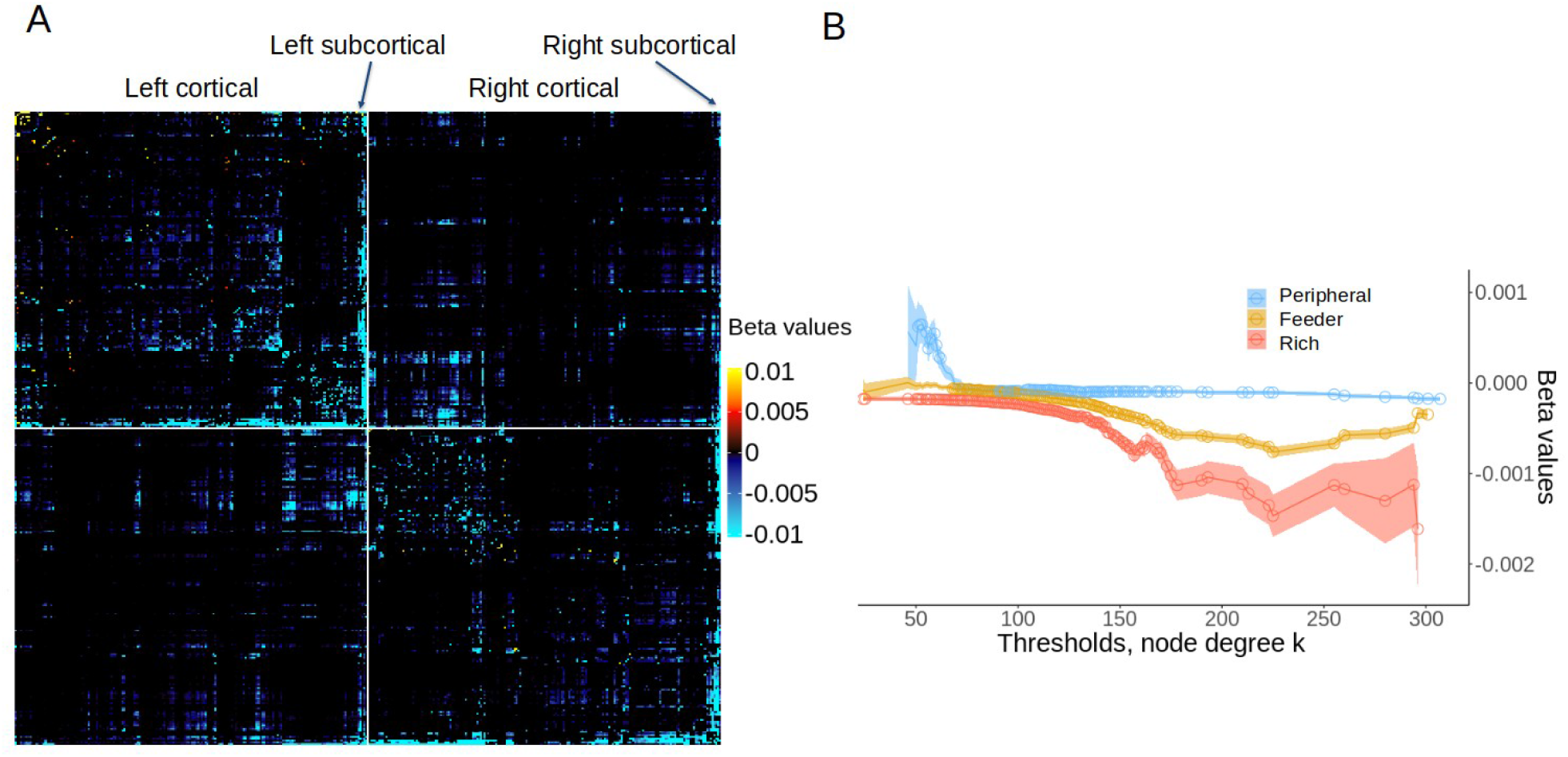
Linear age effects on the whole-brain connectome. (A) Significant (P_TFNBS_ < 0.05) age effects on connectivity matrix at the edge level. The parcellations are divided into 2 parts, left and right hemisphere, and are displayed as: left cortical, left subcortical, right cortical, and right subcortical brain regions. (B) The beta values of averaged connectivity for rich (hub–hub), feeder (hub–nonhub), peripheral (nonhub–nonhub) connections as a function of the thresholds, *k*, used to define hubs. The shaded area indicates one standard error around the mean. Circles represents significant beta-values at a given threshold.

Since many sub-cortical regions have both high degree as well as large age effects (Figure 5), it is possible that the significant spatial correlation between brain maps of the degree and the age effects is driven by the sub-cortical regions. Thus, we repeated the correlation analysis in only cortical regions. The network characterizations of brain networks for only cortical regions are shown in Figure S7. Consistent with the results from the primary analysis, we found that cortical hubs were more susceptible to age-related alterations (r = -0.286, n = 358; Figure S8). Similar, albeit somewhat weaker, results were found for the quadratic effects. A marginally significant spatial correlation between the degrees and the averaged beta-values (P_permutation_ < 0.07, r = -0.173) was found for the analysis that was conducted for only cortical regions (Figure S9).

**Figure 5.**
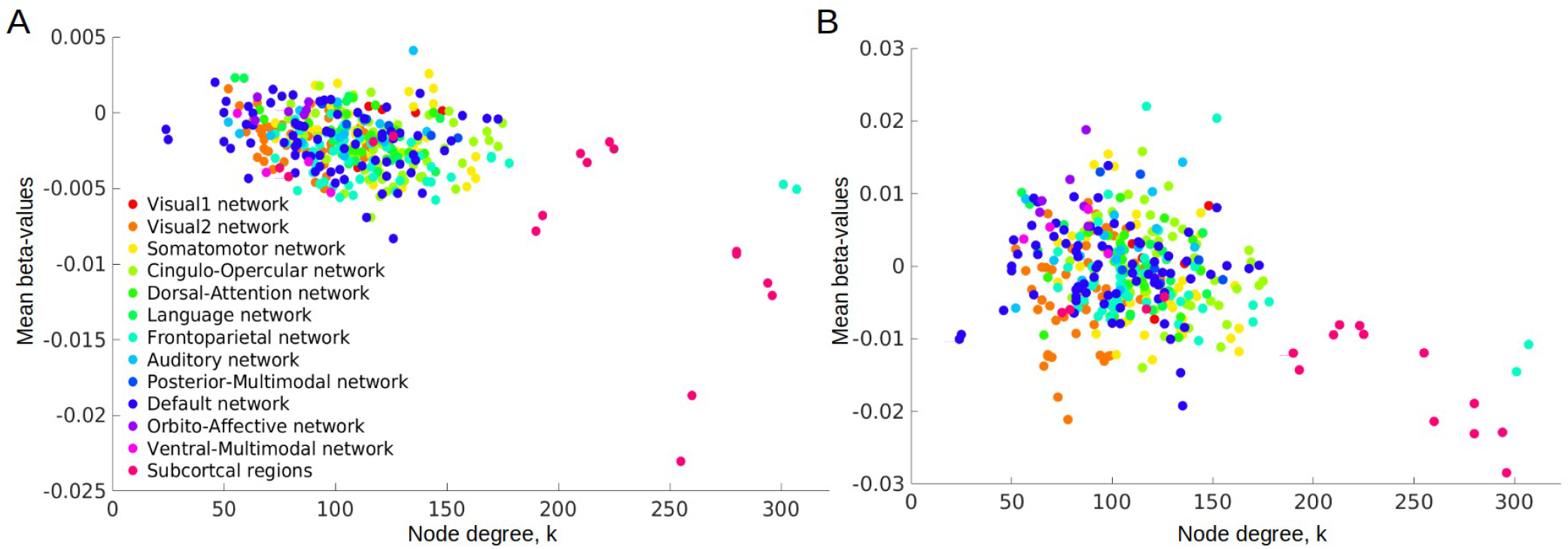
Scatter plots of nodal degrees and the averaged linear (A) and quadratic (B) beta-effects. Colors represent functionally defined networks in cortical regions and sub-cortical brain regions.

In addition to connectivity strength, we found lower clustering coefficient and global efficiency, and higher path length and betweenness centrality in older compared to younger adults (all p-values < 0.00001). Interestingly, the age effects on clustering coefficient, path length and betweenness centrality were larger in hubs compared to non-hub brain regions (p-values < 0.0001, Figure S10).

#### 2.2 Longitudinal changes of the structural connectome

The results showed that across five years, there was a significant decline in structural connectivity (log-transformed FBC) in multiple cortical–cortical connections, as well as cortical–subcortical connections. There were larger changes in intra-hemispheric connections compared to the inter-hemispheric connections. The changes were more pronounced for rich connections compared to feeder and peripheral connections (Figure 6). We also found a weak but significant age effect on the changes of structural connectivity, mainly in rich connections (Figure S11). This result replicates the quadratic age effects and indicates that older adults had larger decline compared to younger adults, and that this age-related difference in decline was most pronounced in rich connections. A correlation analysis showed a significant spatial correspondence between brain maps of the degree and changes in structural connectivity (Figure S12). A similar pattern of results was found for the age effects on 5-year changes in structural connectivity (Figure S13). The longitudinal results generally replicated the main findings of the cross-sectional analysis and further indicates that the decline in structural connectivity in older adults are largest and more concentrated in rich connections. However, compared to the cross-sectional analysis, the longitudinal results demonstrated greater discrepancy in changes between intra- and inter-hemispheric connections. In addition to connectivity strength, we found declined clustering coefficient and global efficiency, and increased path length and betweenness centrality across five years (p-values < 0.00001). The longitudinal changes were greater on betweenness centrality in hubs compared to non-hub brain regions (p-value < 0.01, Figure S14).

**Figure 6.**
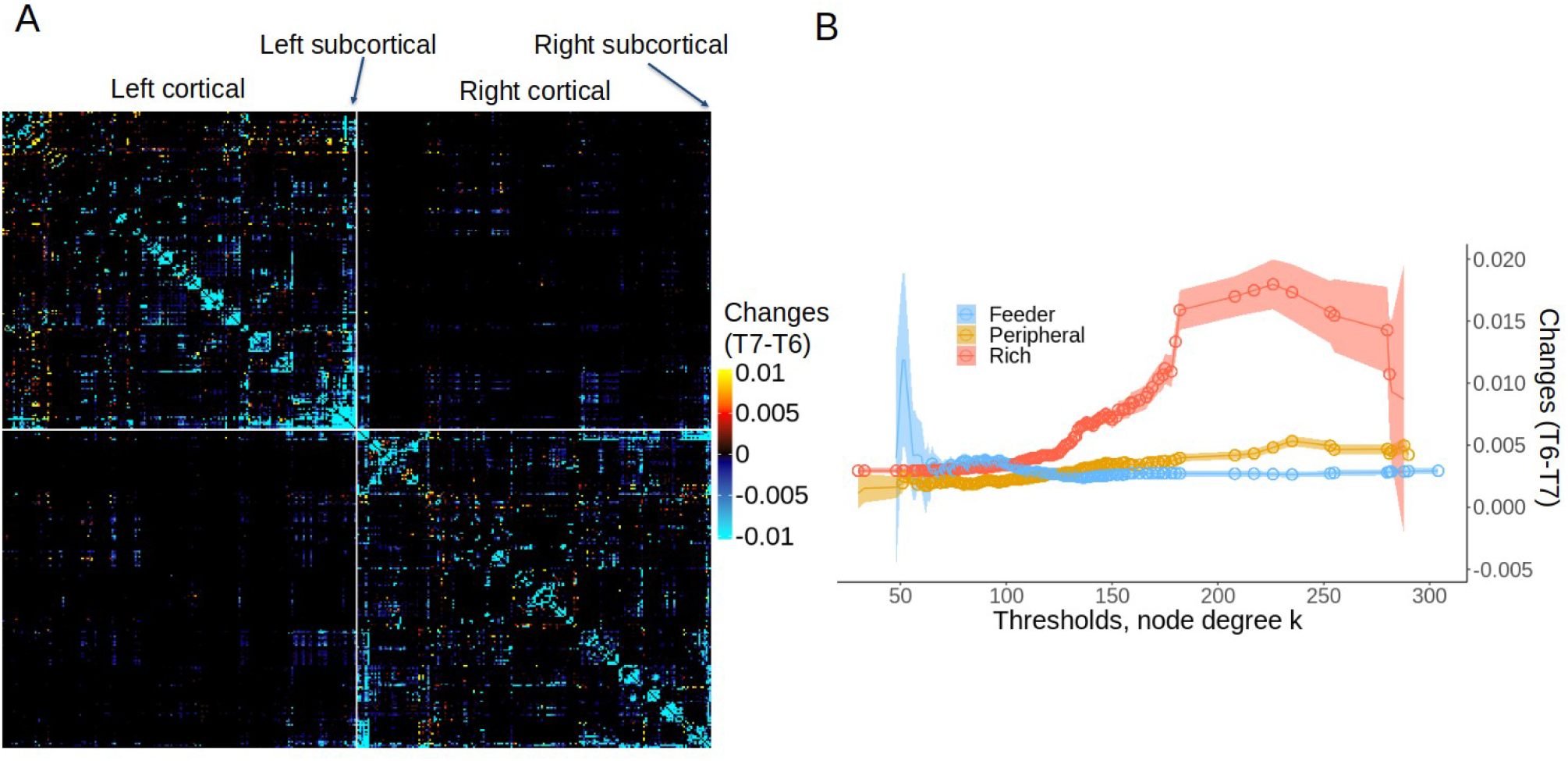
Five year changes on the whole-brain connectome. (A) Significant (P_TFNBS_ < 0.05) changes on connectivity matrix at the edge level. The parcellations are divided into 2 parts, left and right hemisphere, and are ordered as left cortical, left subcortical, right cortical, and right subcortical brain regions. For illustrative purpose, negative changes (T7 – T6) was used in the plot. (B) The averaged connectivity changes for rich (hub–hub), feeder (hub–nonhub), peripheral (nonhub–nonhub) connections as a function of the thresholds, *k*, used to define hubs. The shaded area indicates 1 standard error around the mean. Circles represents significant changes at a given threshold.

### 3. The relation between structural connectivity and processing speed

TFNBS analyses showed significant associations between multiple cortical/subcortical structural connections and processing speed. The associations were found in the connections between somatosensory and motor areas, inferior frontal cortex, inferior and superior parietal cortex, lateral temporal cortex, the temporo-parieto-occipital junction, posterior opercular area, auditory areas, and a subcortical region in ventral diencephalon (Figure 7). A majority (13 out or 14) of the connections that were significantly (P_TFNBS_ < 0.05) associated with speed, also had significant (P_TFNBS_ < 0.05) linear or quadratic age effects.

**Figure 7.**
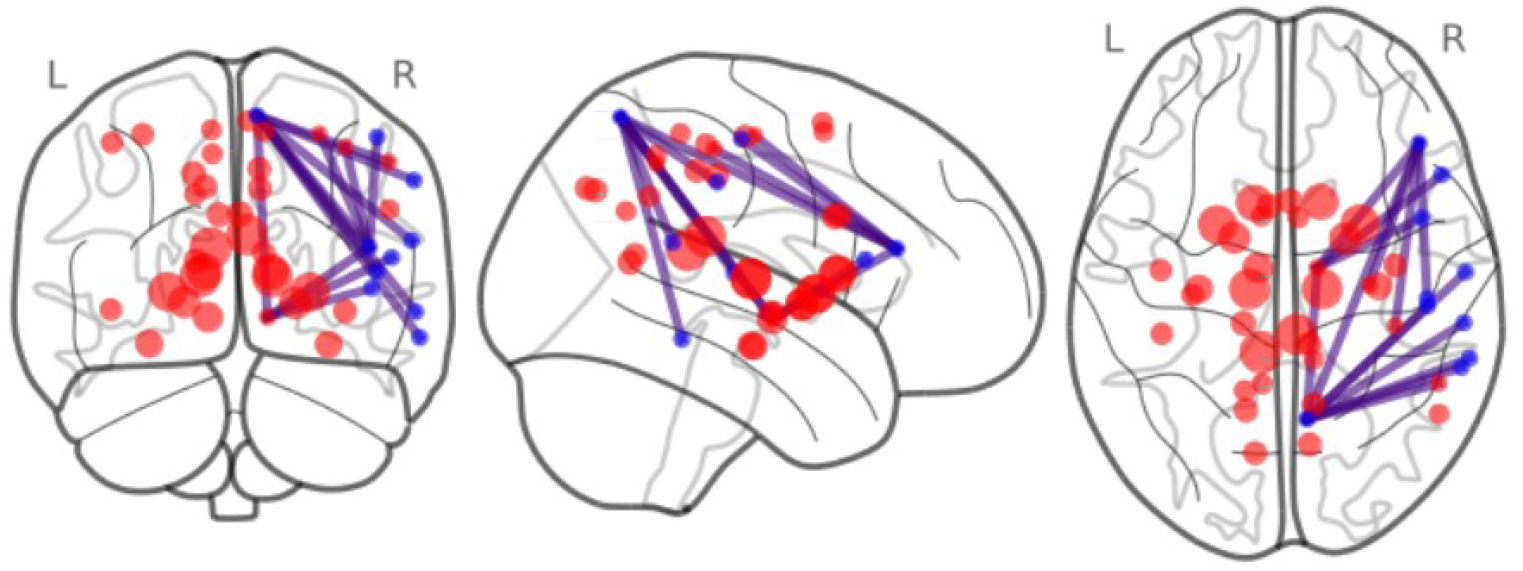
Connections (purple) that showed significant associations between connectivity strength and processing speed; red circles represent hubs defined by 1 SD above the mean degree; blue circles represent non-hubs; the size indicates the degree of each node.

We tested if the connections that were significantly associated with processing speed were more concentrated in hub–hub connections. The median of degrees for the brain regions of 14 significant connections is 197, which is significantly larger than expected by chance (P_permutation_ < 0.0001, Figure S15A), indicating that brain regions with larger degrees were also the ones that were significantly related to processing speed. We also found that the mean beta-value that was associated with processing speed within the rich connections was significantly larger than the null distribution (P_permutation_ < 0.001), whereas the mean beta-value within the peripheral connections is significantly lower than the null distribution (p < 0.001, Figure S15B). However, it should be noted that this analysis was conducted on the whole network, not only within the 14 connections that showed significant associations with speed. These results suggest that the largest associations between structural connectivity and processing speed are concentrated in rich connections within a brain network. The significant mediation effect (p < 0.0001) showed that 16.7% of the total effect between age and processing speed can be explained by differences in connectivity strength in hub-like regions.

Using longitudinal data, we also tested the change-change associations between connectivity and processing speed from baseline to follow-up using TFNBS controlling for age and sex (△ *Speed* ∼ *Age*+ *Sex* +log (△ *FBC*)). However, no significant change-change correlations was found between FBC and processing speed across five years.

### 4. Validation

Instead of degree, the analysis using hubness to define the level of central/influential of brain regions in the network replicated the finding that the age effects were largest in rich connections, compared to feeder or peripheral connections (Figure S16). We also tested if the associations between brain maps of the degrees and the beta-values of age/speed are significant after controlling for differences in brain volumes. Indeed, we were able to replicate our finding using different parcellations, connectome thresholds, or the definitions of connectivity strength (Figure S17). These supplementary analyses show that the pattern of results replicate well and are not influenced by differences in hub definition or parameters used in the analyses.

## Discussion

Three main findings were obtained in the current study. First, although the anatomical distributions of hubs in the structural brain network are relatively stable across age, the connectivity strength among hub-like regions are more susceptible to age-related insults compared to non-hub connections. These findings were supported by both cross-sectional and longitudinal analyses. Second, the associations between connectivity strength and processing speed were larger in hub compared to non-hub connections. Third, age-related differences in connectivity strength within hub connections accounts for lower processing speed in older adults. These findings provide evidence that hubs in the structural brain network are more affected by normal aging, and that alterations in hub connectivity partly explain age-related deficits in processing speed in older adults.

Using a novel measure of connectivity, FBC, we demonstrated that the human structural brain network has a rich-club organization with a long-tailed degree distribution, the mean distance of connections increases as a function of increasing degree, and that the degrees are related to multiple measures of regional brain metabolism. These topographical characteristics are consistent with previous studies (Arnatkeviciute et al., 2021; Crossley et al., 2014; Xu et al., 2022), indicating that the brain network is not random but rather a scale-free network with a subset of hubs having long-range connections and high energy cost.

The current results demonstrated similar anatomical locations of hubs across the adult life-span, but a less global youth-like degree distribution with advancing age. We also show that the clustering coefficient, global efficiency, path length, and betweenness centrality differed between younger and older adults, especially within hub connections. These findings indicate a relative age-invariant spatial distribution of structural hubs in older adults, but also alterations of the whole-brain topological organization in older adults. The cross-sectional TFNBS analysis demonstrated widespread age effects on structural connectivity strength, as indicated by lower FBC with increasing age. Similar results were observed in the longitudinal analysis, showing that FBC declined globally across five years in older adults. These results corroborate previous findings of age effects using multiple indicators of WM microstructure, including WM integrity (e.g., Sexton et al., 2014), WM fiber density (Choy et al., 2020; Kelley et al., 2021), and number of streamline counts (Coelho et al., 2021). Together with the present results, these findings indicate robust and replicable effects of age on WM microstructure across measurements and using different biophysical models of water diffusion. The finding of lower structural connectivity strength in older compared to younger adults support previous work showing that the degree to which structural brain networks rely on geometric wiring rules is weakened with advancing age (Betzel et al., 2016; Zuo et al., 2017). This finding may also indicate less structural constraints on functional brain organization with advancing age, which might lead to disassociation of the topographical organizations between structural and function connectivity in older adults (Zamani Esfahlani et al., 2022).

Importantly, the current results extend previous observations by demonstrating that the age effects on structural connectivity are heterogeneous, with more pronounced effects in hub compared to peripheral connections. This pattern was consistent both across thresholds and irrespective of analytical choice for defining hubs. Since hubs generally have longer-range connections, they are particularly important for connecting brain sub-networks (Van Den Heuvel et al., 2012). Weakened connections in hub regions might therefore lead to a more differentiation of structural organization characterized by decreased integration and increased segregation of the network shown as demonstrated in previous studies (Coelho et al. 2020). Moreover, computational work has shown that the complex scale-free network has extremely high tolerance for random errors (Albert et al., 2000), but demonstrates rapid deterioration of global efficiency when hubs are attacked by modeled pathological damage (Crossley et al., 2014). Our findings that hub connections are differentially more susceptible to the aging process might explain lower global network efficiency in older compared to younger adults.

While structural connectivity patterns from cross-sectional and longitudinal analyses largely converged, some differences were also identified. The longitudinal results showed larger effects of time on intra-compared to inter-hemispheric connections. One possible reason for this difference is that, compared to cross-sectional data, longitudinal analyses might be less influenced by confounding factors, such as cohort effects (e.g., education and nutrition). Additional longitudinal studies are clearly needed to characterize age-related changes in intra- and inter-hemispheric network connectivity.

It is currently not clear why the effect of age is selectively larger in hub-like regions compared to peripheral connections. There might be several potential explanations. First, since hub connections are generally longer and have been associated with higher blood flow and/or metabolic rate, it is possible that oxidative stress and energy depletion induced by the accumulation of RONCS during aging have a particular impact on hubs because of their higher demands for energy and vulnerability to oxidative stress. Second, since older adults have lower endogenous dopamine (DA) level and DA D1/D2 receptor density compared to younger adults (for a review, see Karrer et al., 2017), a lower level of dopamine could result in altered WM connectivity. In support of this view, some studies have linked DA function with WM integrity (Li et al., 2022). Animal work has also shown that DA-depleted rats have decreased length of dendrites, as well as altered neuronal density (Kalsbeek et al., 1989, 1987; Wang and Deutch, 2008). Moreover, there are well-established links between measures of dopamine D1 (Johansson et al., 2022), D2 (Karalija et al., 2019), and DA transporter (Rieckmann et al., 2016) and WM lesions. In the current study, we note that brain regions associated with the high dopamine D1/D2 receptor availability, such as the basal ganglia and the prefrontal cortex also have higher degrees compared to other brain regions. It is therefore possible that age-related decline of DA in these regions contribute to deterioration of the WM fibers, which may partly lead to more deleterious age effect on connections of these nodes.

The current results also suggests that hubs may be particularly critical for human cognition. Consistent with previous studies, we found that structural connectivity was associated with processing speed (Zimmermann et al., 2018). The current results add new information about the relation between cognitive performance and WM integrity by demonstrating that the relation was particularly linked to hub connections. Indeed, hubs are critical for information transformation among long-distance connections and increases global network efficiency. cognitive performance has been related to global efficiency in both structural (Pineda-Pardo et al., 2016) and resting-state functional (Gießing et al., 2013; Van Den Heuvel et al., 2009) brain networks. Our results reveal an important role of structural hubs in tasks that require efficient information processing, such as in processing speed. Moreover, processing speed has been related to cognitive performance of many task domains. Slower processing speed has been suggested as an underlying cause of accelerated cognitive impairment in older adults (Finkel et al., 2005). Thus, our results extend previous findings by indicating that lower connectivity strength within hub connections may be one of the neural mechanisms of cognitive decline, especially in processing speed, among older adults.

While we believe that the current study has several strengths, including a relatively large sample size with a wide age range, and the availability of both cross-sectional and longitudinal data, a few limitations should be highlighted. First, in this study we focused on degrees and hubs, which are the key characters in graph theory, as indicators of structural connectivity. However, there are many other topological properties, such as small-worldness and modularity, which reflect different aspects of connectomic organization. Future studies are clearly needed to explore how these properties relate to each other, how they are affected by adult aging and their association with various cognitive functions. We believe that a more systematic characterization on multiple measures of connectome organization could provide a deeper understanding about the complex human network and how age might influence those properties of the network. Second, the organization and characteristics of the structural and functional brain networks are similar but not entirely identical. Here we specifically examined structural brain network organization, and therefore lack information on how these results translate to the functional dynamics of hub organization. Future studies that include multimodal assessment of the human connectome are required to explore how such structure-function associations related to cognitive functioning in older adults.

## Conclusion

By combining network-based and graph-theoretical analyses, the current study extends previous findings by showing age-related alterations in structural connectivity strength, along with relative unchanged spatial brain network distributions of hubs with advancing age. Importantly, results from both cross-sectional and longitudinal analyses demonstrate that the connections between hubs are more strongly influenced by aging compared to peripheral regions, which might explain the alterations of whole-brain topological organizations, such as lower global efficiency in older adults. In addition, we found that the hub connections mediated the effect of age on processing speed, suggesting that weaker hub connections are behaviorally detrimental and at least partly explains lower processing speed in older adults.

## Acknowledgments

This work was supported by the Swedish Research Council [421-2013-1039] and Hjärnfonden [F02014-0224] to J.P. Grants from the Swedish Research Council supported the BETULA Project.

