## Supplementary figures for "Hub architecture of the human structural connectome: Links to aging and processing speed"

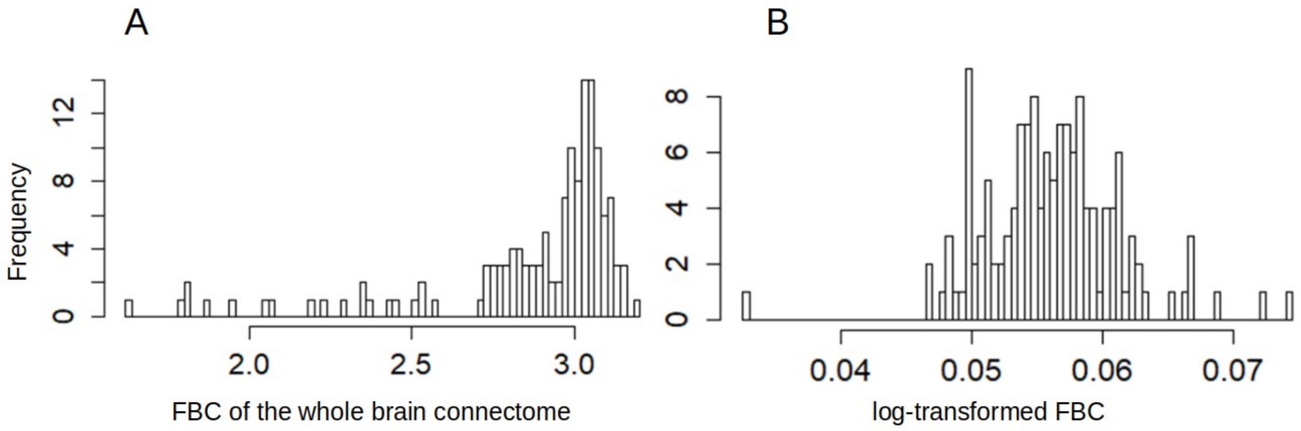


Figure S1. Distributions of fiber bundle capacity (FBC) and log-transformed FBC of the whole connectome. The connectome is based on the average of the baseline sample (N=137).

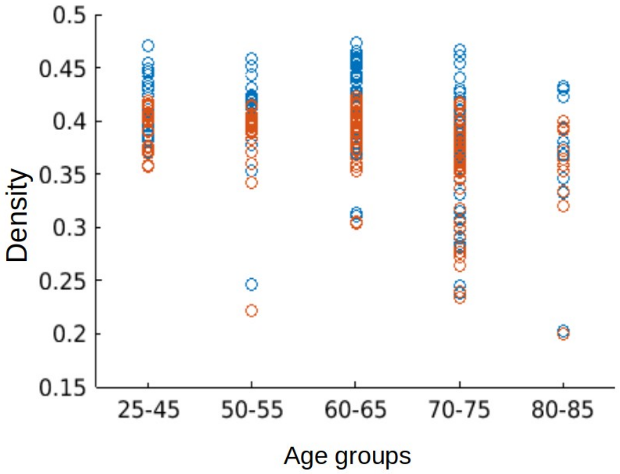


Figure S2. Distributions of connection densities of the original connectome (blue) and the connectome after thresholding (red) across age groups.

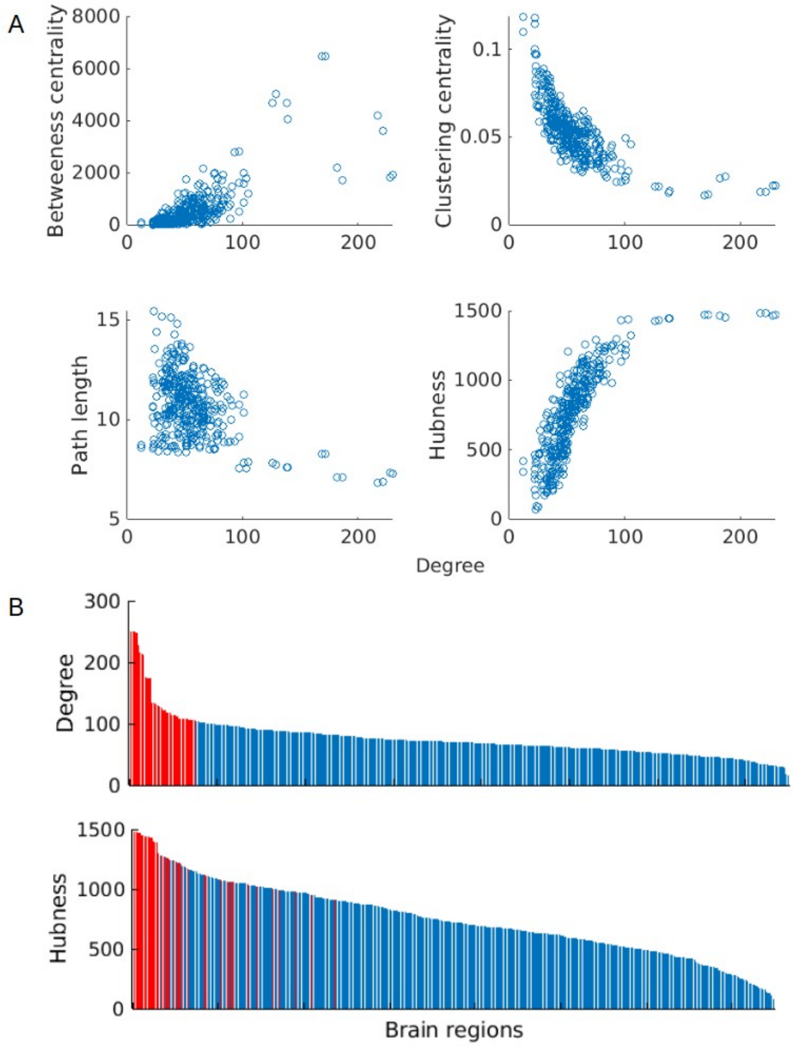


Figure S3. Degree of the mean connectome was highly correlated with hubness and with other measurements of centrality, i.e., path length, betweenness centrality, and clustering-coefficient (A). Hubs defined by degree are consistently ranked for the measure of hubness (B)

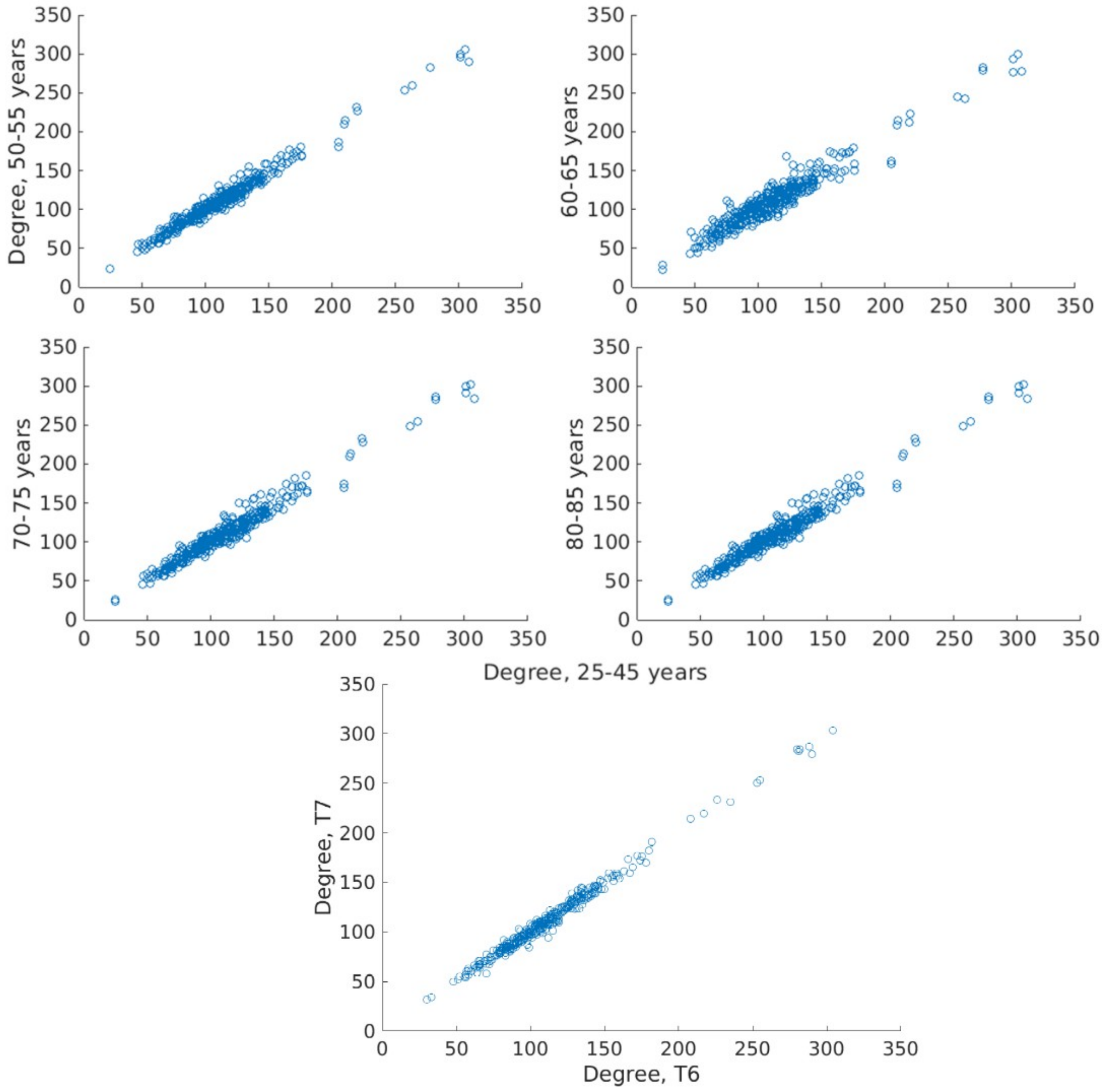


Figure S4. The distributions of degrees across age groups and time points are highly correlated.

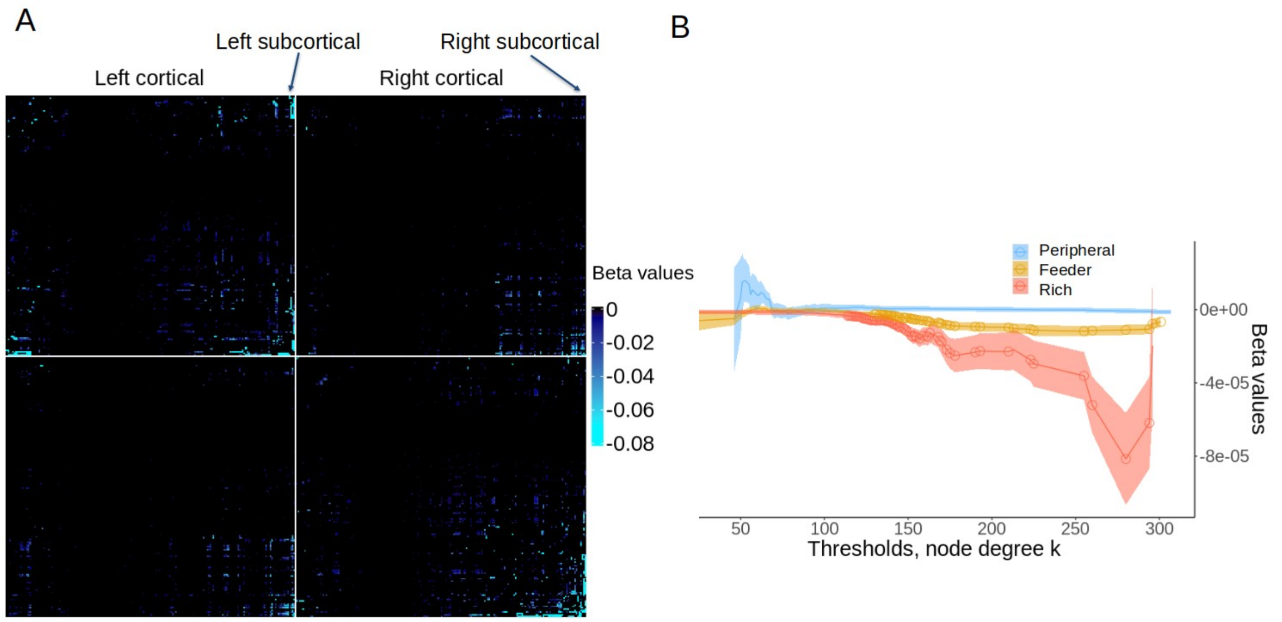


Figure S5. Quadratic age effects on the whole-brain connectome. (A) Significant ($P_{TFNBS}$ <0.05) age effects on the connectivity matrix at the edge level. The parcellations are divided into 2 parts, left and right hemisphere, and are ordered as left cortical, left subcortical, right cortical, and right subcortical brain regions. Only negative quadratic age effects are significant, and positive beta-values are therefore not displayed. (B) The beta values of averaged connectivity for rich (hub–hub), feeder (hub–non-hub), peripheral (non-hub–non-hub) connections as a function of the thresholds, *k*, used to define hubs. Shaded area is one standard error around mean. Circles represents significant beta-values at a given threshold.

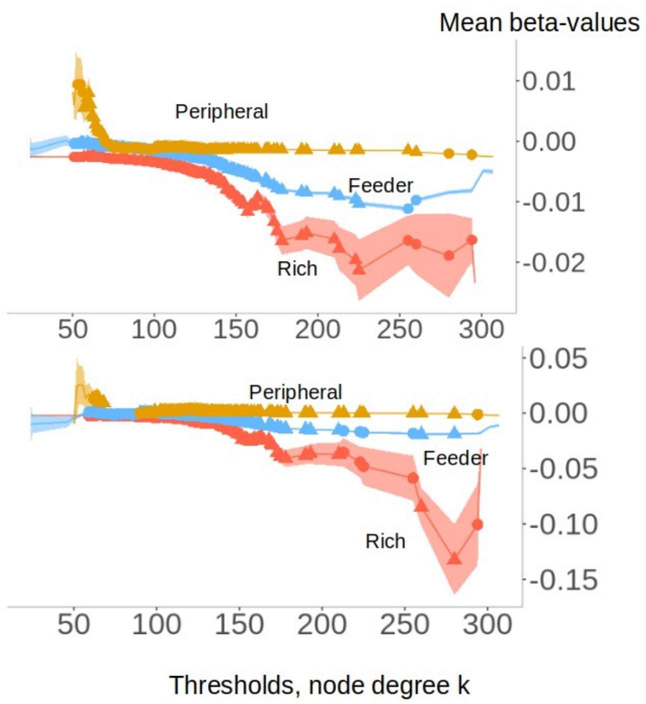


Figure S6. Mean linear (top) and quadratic (bottom) beta values were larger for rich compared to feeder (linear: $p < 0.005$ for $175 > k > 46$; quadratic: $p < 0.005$ for $168 > k > 85$) and peripheral connections (linear: $p < 0.0001$ for $k > 102$; $p < 0.005$ for $k > 103$) across degrees.

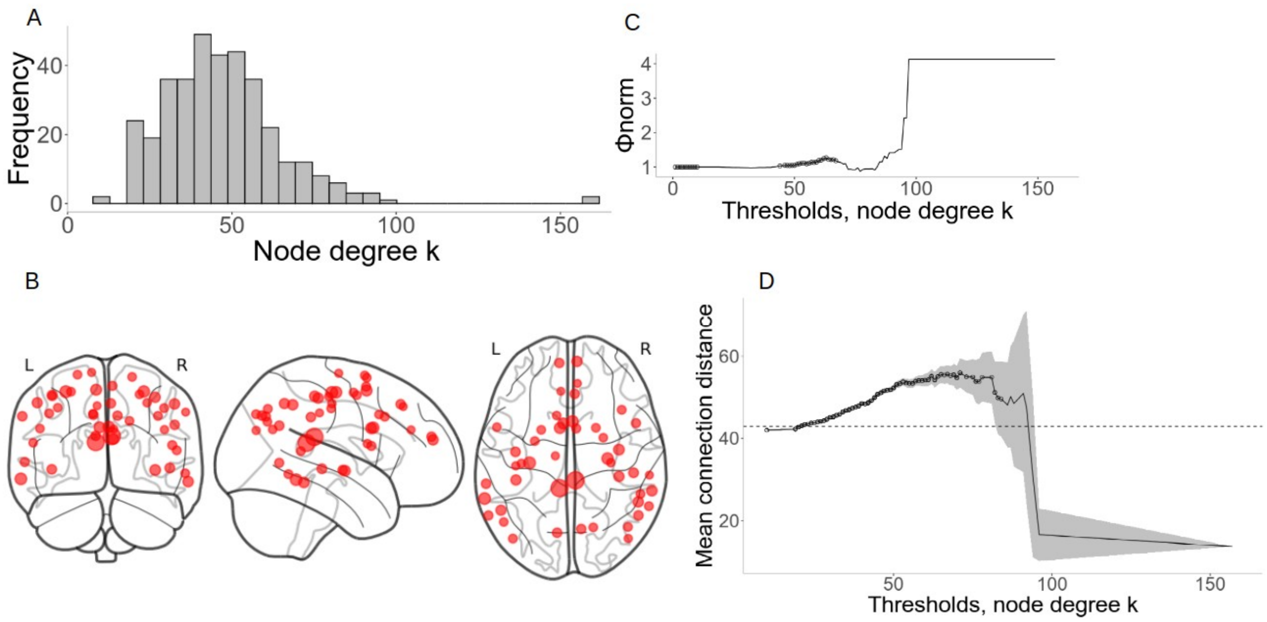


Figure S7. Network characteristics of cortical brain regions. (A) The degree distribution across nodes. (B) The anatomical locations of hubs defined by 1 SD above the mean degree; the size indicated the degree of each node. (C) The normalized rich-club coefficient φ norm increases as a function of degree threshold (*k*) used to define hubs. Black circles indicate significant ($P_{permutation} < 0.05$) φnorm compared to 1000 degree-retained random networks. (D) Mean connection distance among rich club (hub-hub) connections increases as a function of degree threshold (*k*) used to define hubs. The shaded area depicts two standard errors around the mean. Black circles indicate significant larger distance among rich club connections compared to averaged feeder and peripheral connections. The dashed line shows the average connection distance across all network links.

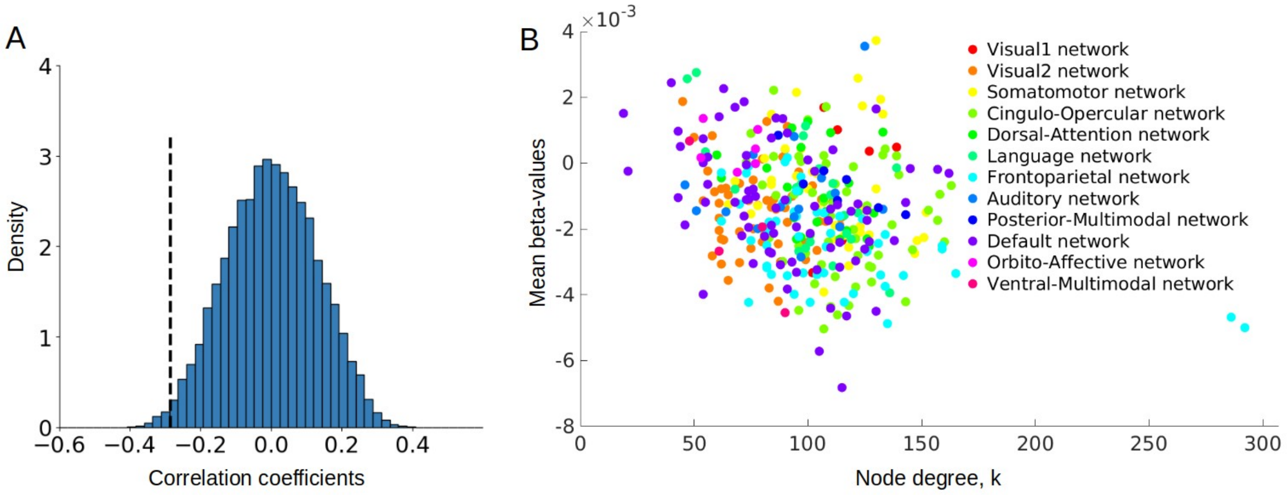


Figure S8. (A) The distribution of Pearson's correlations between degree and 10.000 SA-preserved surrogate brain maps of the linear age effects for only the connections in cortical regions. The dashed lines represent the true correlations based on our data. (B) Scatter plot of nodal degree and the averaged beta-effects. Colors represent functionally defined networks in cortical regions.

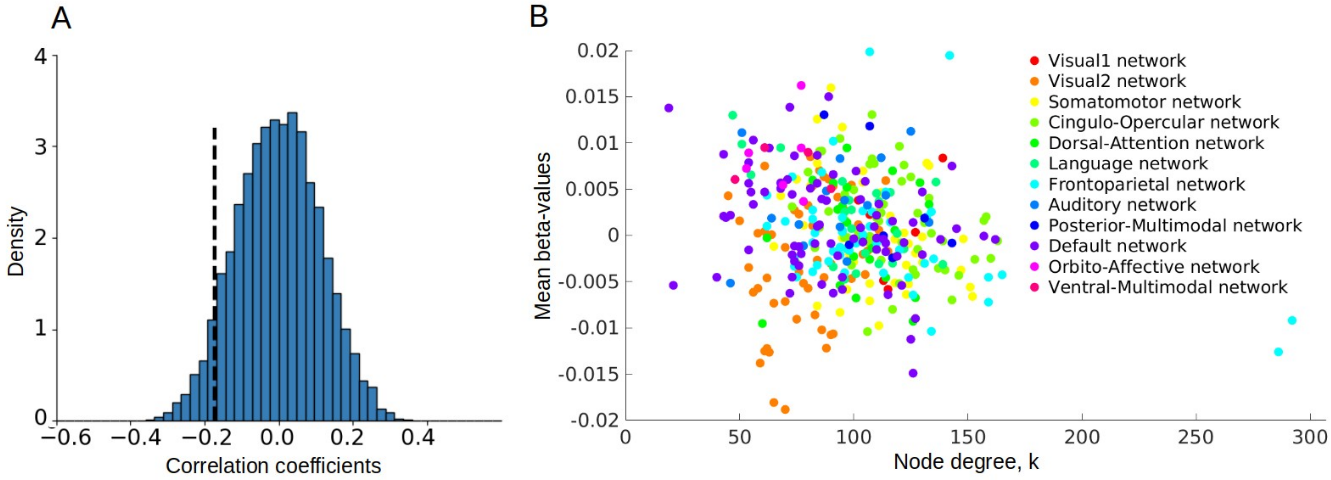


Figure S9. (A) The distribution of Pearson's correlations between degree and 10.000 SA-preserved surrogate brain maps of the quadratic age effects for only the connections in cortical regions. The dashed lines represent the true correlations based on our data. (B) Scatter plot of nodal degree and the averaged beta-effects. Colors represent functionally defined networks in cortical regions.

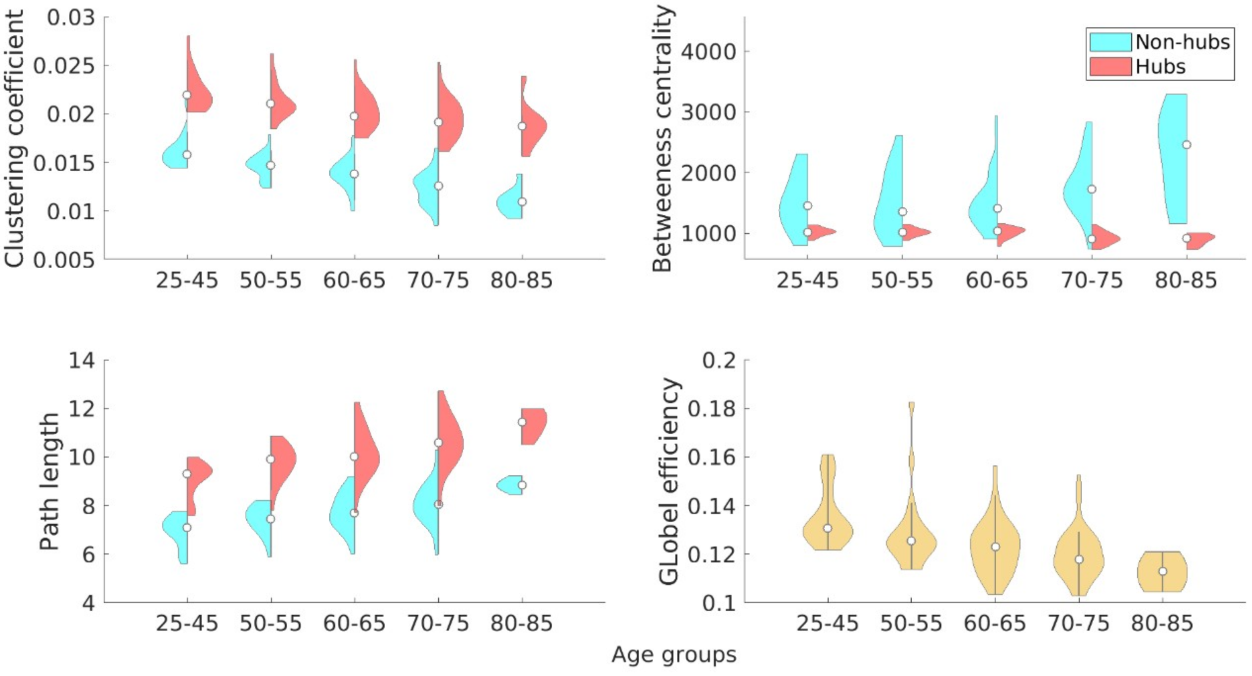


Figure S10. Clustering coefficient, global efficiency, path length, and betweenness centrality of hub/non-hub brain regions across age groups. Hubs are defined as the brain regions with top 5% highest degrees.

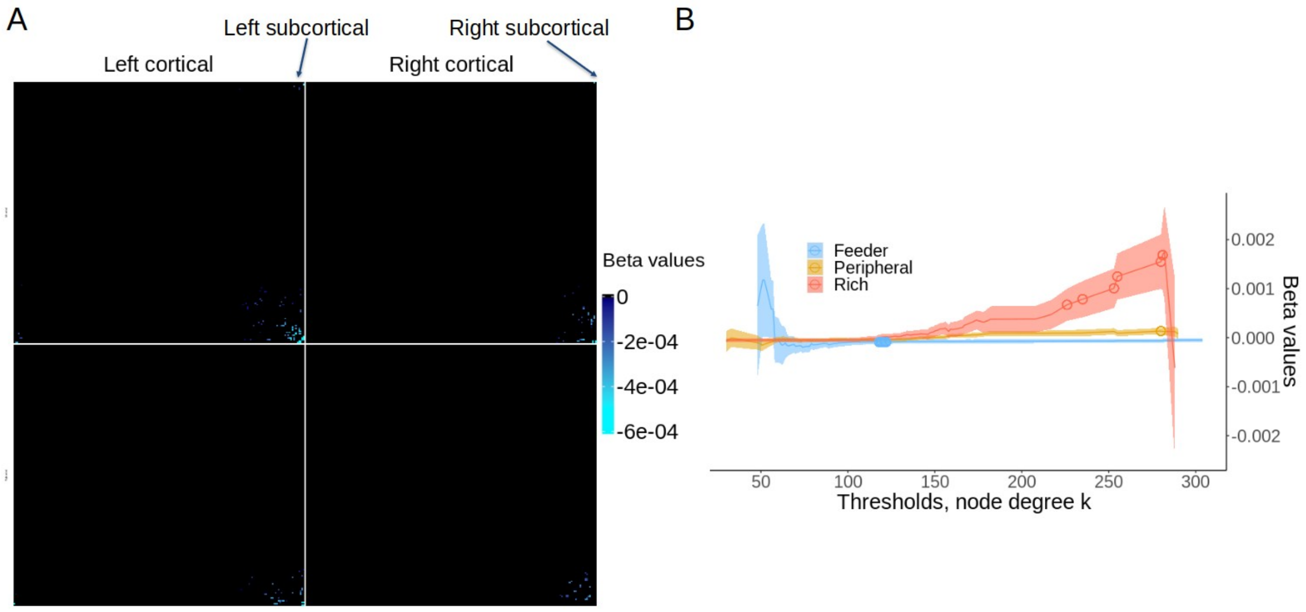


Figure S11. The age effects on 5-year changes in the whole-brain connectome. (A) Significant ($P_{TFNBS} < 0.05$) age effects on the changes of connectivity matrix at the edge level. The parcellations are divided into 2 parts, left and right hemisphere, and are ordered as left cortical, left subcortical, right cortical, and right subcortical brain regions. For illustrative purpose, negative changes, i.e., T7-T6, was used in this plot. (B) The averaged beta-values of age on the connectivity changes for rich (hub–hub), feeder (hub–non-hub), peripheral (non-hub–non-hub) connections as a function of the thresholds, *k*, used to define hubs. The shaded area represents 1 standard error around the mean. Circles represents significant age effects on the changes at a given threshold.

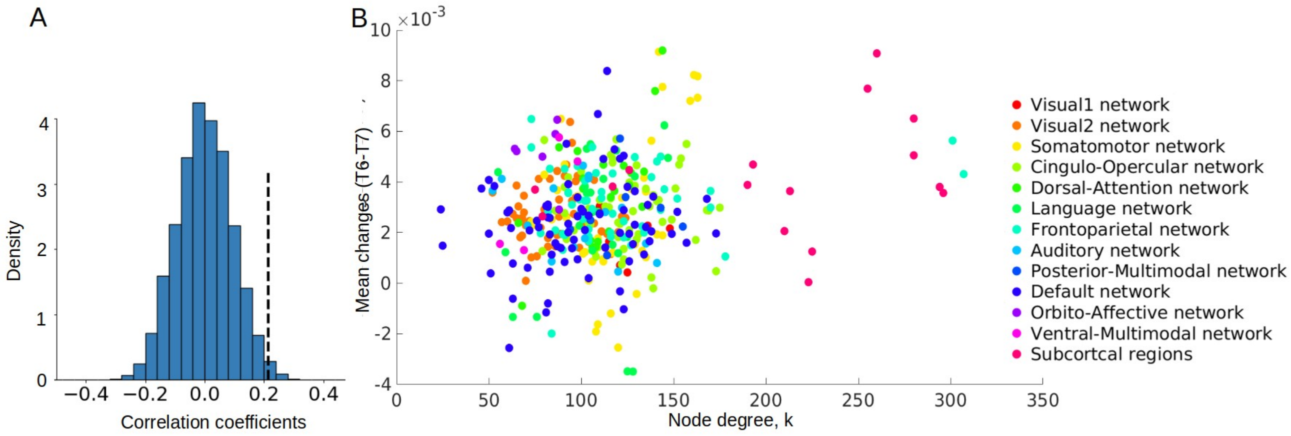


Figure S12. (A) The distribution of Pearson's correlations between degree and SA-preserved surrogate brain maps of the changes in log-transformed FBC. The dashed lines represent the true correlations based on our data. (B) Scatter plot of nodal degree and the averaged changes from T6 to T7. Colors represent functionally defined networks in cortical and subcortical regions.

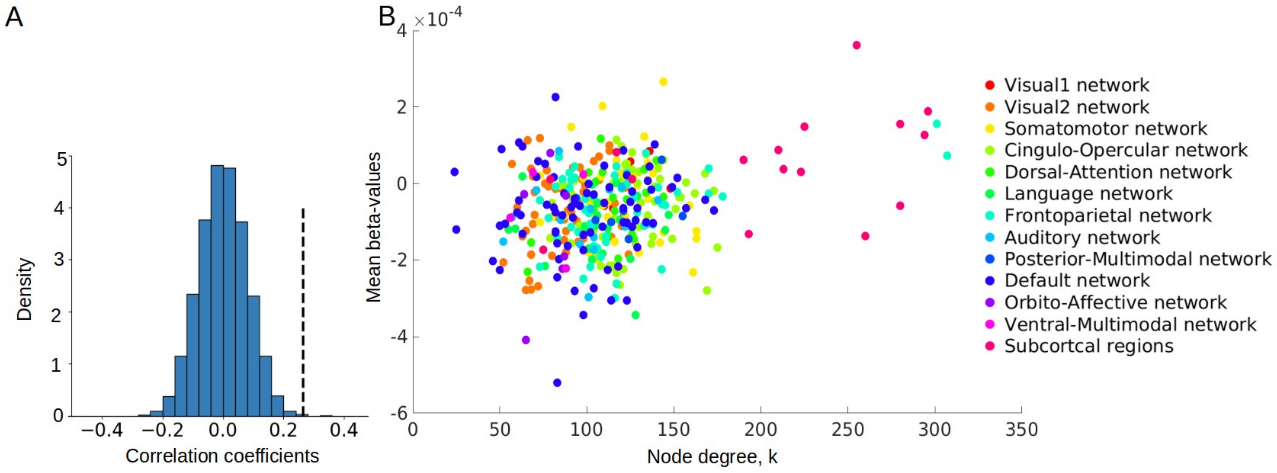


Figure S13. (A) The distribution of Pearson's correlations between degrees and SA-preserved surrogate brain maps of the age effects on the changes in log-transformed FBC. The dashed lines represent the true correlations based on our data. (B) Scatter plot of nodal degree and the averaged beta-values of age effects on the changes from T6 to T7. Colors represent functionally defined networks in cortical and subcortical regions.

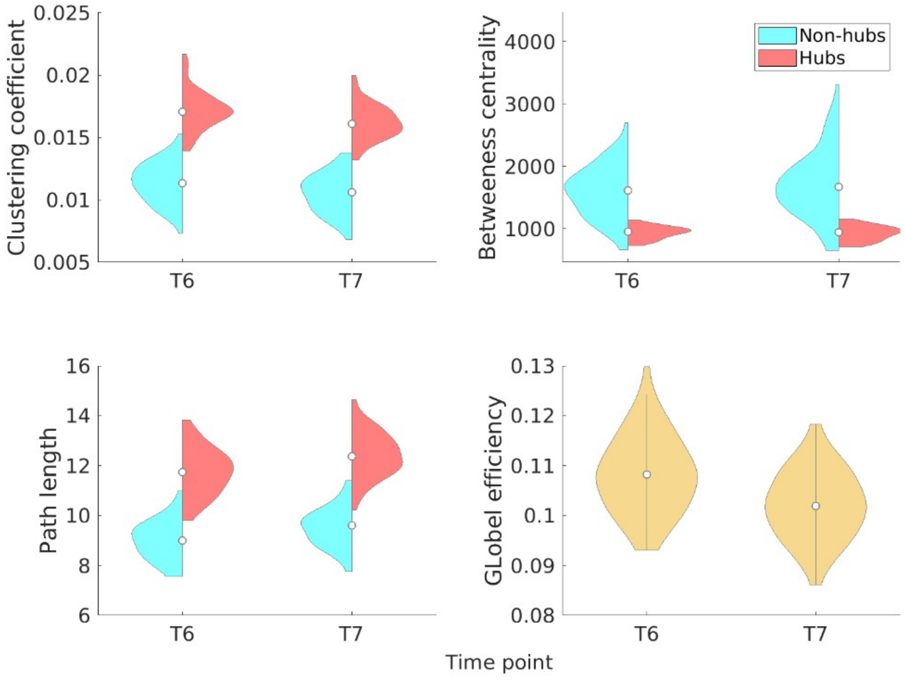


Figure S14. Clustering coefficient, global efficiency, path length, and betweenness centrality of hub/non-hub brain regions across time points. Hubs are defined as the brain regions with top 5% highest degrees.

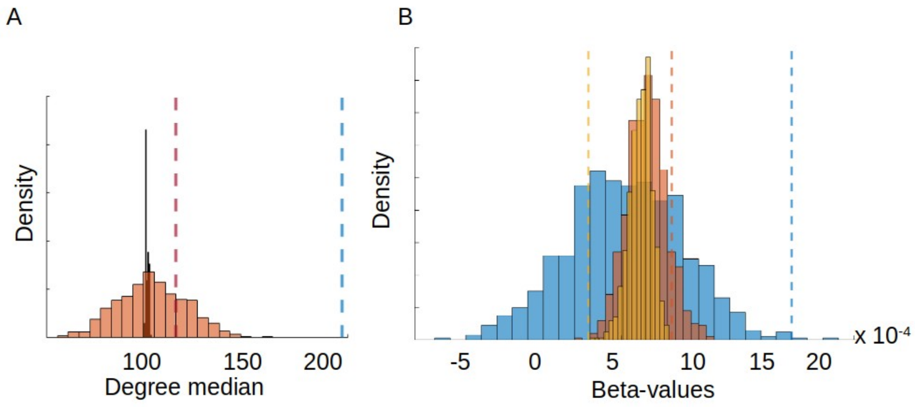


Figure S15. (A) The distribution of median degree for the connections that were associated with processing speed (orange), and for the remaining connections in the network (red). The distribution is based on 10.000 SA-preserved surrogate maps of degrees. Dashed lines represent the median degree for the connections that were associated with speed (blue), and for the rest connections in the network (red) based on our true data. (B) The distribution of mean beta-values of the associations with speed for rich (blue), feeder (red), and peripheral (yellow) connections. The distribution is based on 10.000 SA-preserved surrogate maps of degrees. Dashed lines represent the averaged beta-values of speed for rich (blue), feeder (red), and peripheral (yellow) connections base on our data.

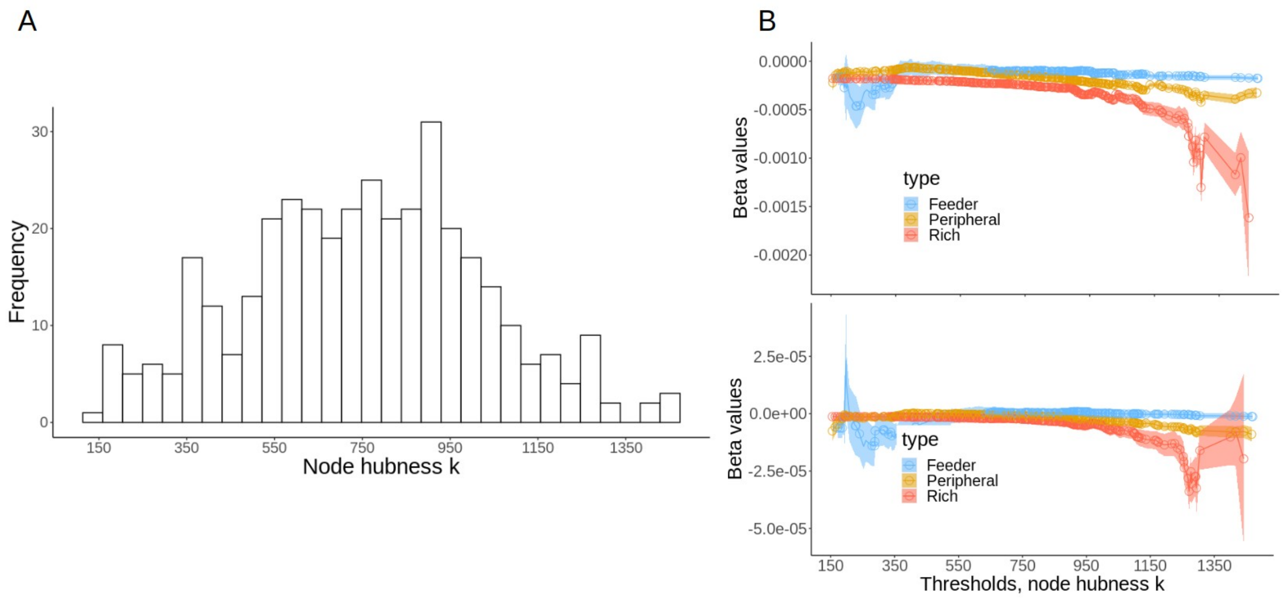


Figure S16. (A) The distribution of hubness across nodes. (B) The beta values of the linear (top) and quadratic (bottom) on the averaged connectivity for rich, feeder, peripheral connections as a function of hubness, used to define hubs. Shaded area is one standard error around mean. Circles represents significant beta-values at a given threshold.

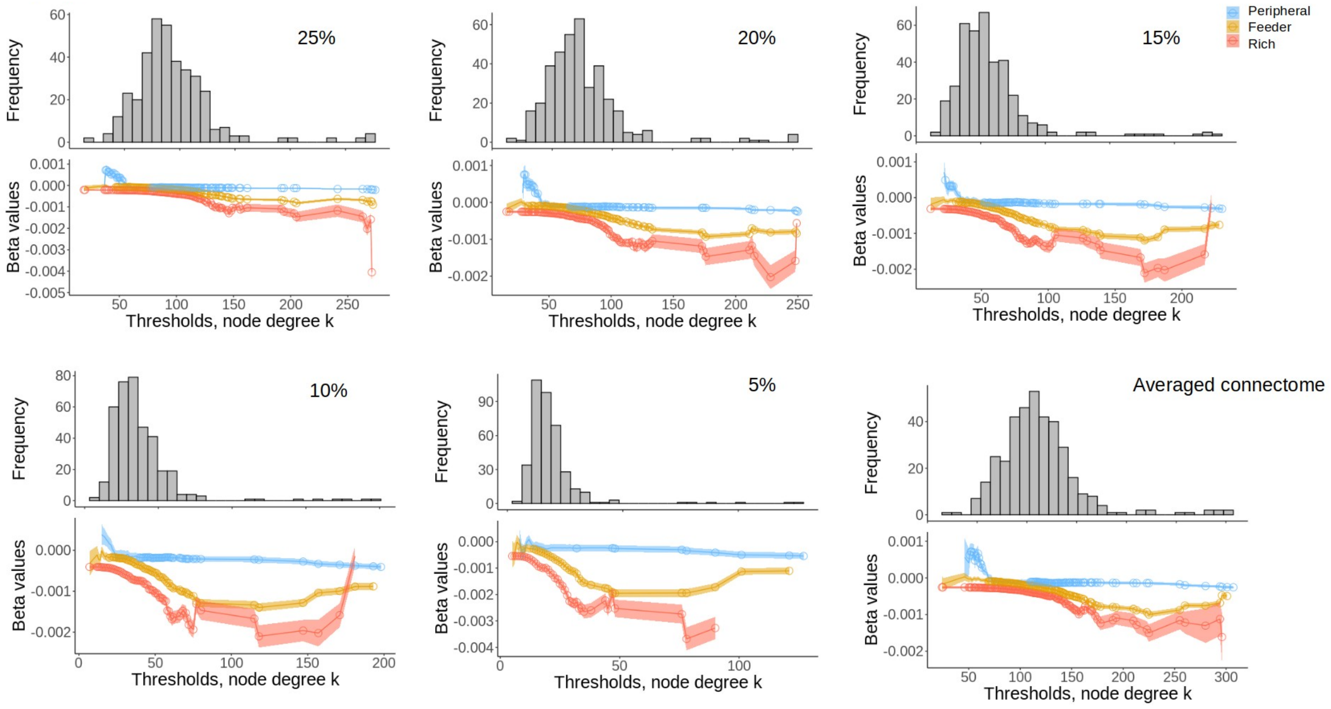


Figure S17. Validation analysis show repeated patterns of the main findings based on different densities of the adjacent matrix (thresholding ii based on τ = 5 %, 10 %, 15 %, 20 %, 25%, and on the mean connectome for all the density thresholds including τ = 30 % in the main analysis). In each subplot: (Top) the long-tailed degree distribution indicates the presence of a small number of highly connected hubs. (Bottom) the beta values of averaged connectivity for rich (red), feeder (yellow), and peripheral (blue) connections as a function of the thresholds, *k*, used to define hubs. The shaded area indicates one standard error around the mean. Circles represents significant beta-values at a given threshold.
